# A hold-and-feed mechanism drives directional DNA loop extrusion by condensin

**DOI:** 10.1101/2021.10.29.466147

**Authors:** Indra A. Shaltiel, Sumanjit Datta, Léa Lecomte, Markus Hassler, Marc Kschonsak, Sol Bravo, Catherine Stober, Sebastian Eustermann, Christian H. Haering

## Abstract

SMC protein complexes structure genomes by extruding DNA loops, but the molecular mechanism that underlies their activity has remained unknown. We show that the active condensin complex entraps the bases of a DNA loop in two separate chambers. Single-molecule and cryo-electron microscopy provide evidence for a power-stroke movement at the first chamber that feeds DNA into the SMC-kleisin ring upon ATP binding, while the second chamber holds on upstream of the same DNA double helix. Unlocking the strict separation of ‘motor’ and ‘anchor’ chambers turns condensin from a one-sided into a bidirectional DNA loop extruder. We conclude that the orientation of two topologically bound DNA segments during the course of the SMC reaction cycle determines the directionality of DNA loop extrusion.

## Main text

Members of the SMC (structural maintenance of chromosomes) family of protein complexes have recently emerged as a new class of molecular motors that perform mechanical work on DNA (*1, 2*). In eukaryotes, the cohesin SMC complex delimits large intra-chromosomal loops that are thought to control gene expression during interphase (*3*) and the condensin SMC complex creates arrays of loops that form the structural basis of rod-shaped mitotic chromosomes (*4, 5*). Single-molecule experiments have demonstrated that both complexes can create and processively enlarge DNA loops over tens of kilo-base pairs (kbp) *in vitro* (*6-9*). In these experiments, condensin primarily reeled in DNA from only one side, while cohesin incorporated DNA into the growing loop from both sides.

The molecular mechanism by which these motors couple adenosine triphosphate (ATP) hydrolysis to DNA loop expansion remains unresolved and faces the challenge that it must account for both symmetric and asymmetric loop extrusion by architecturally similar protein complexes. Both complexes are built around a heterodimer of SMC protein subunits that dimerize at a ‘hinge’ domain located at the end of ∼40-nm-long anti-parallel coiled coils (**Fig. 1A**). Sandwiching of two ATP molecules creates a temporary second dimerization interface between ‘head’ domains at the other end of the coils, which are flexibly connected by a largely unstructured kleisin subunit even in the absence of nucleotide. The central region of the kleisin is bound by two subunits that are composed of consecutive HEAT (Huntingtin, EF3A, PP2A, TOR) repeat motifs (*10, 11*) and have the capacity to interact with DNA and the SMC ATPase heads (*12-18*).

**Fig. 1.**
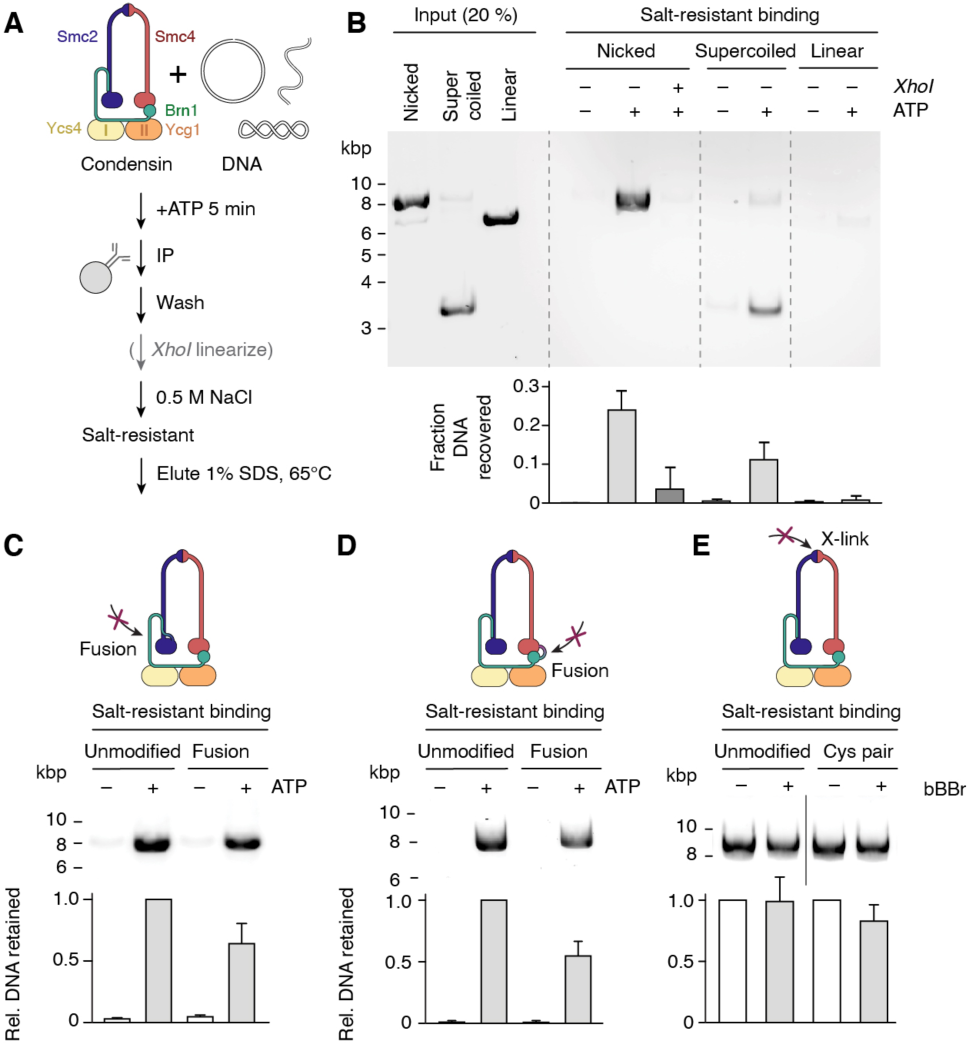
ATP-dependent topological DNA loading of condensin without SMC-kleisin ring opening. **(A**) Schematic of the *in-vitro* DNA loading assay. (**B**) Distinct DNA topoisomers bound to condensin after 0.5 M NaCl washing were eluted with 1 % SDS, resolved by agarose gel electrophoresis and quantitated after ethidium bromide staining (mean ± s.d., *n* = 4). (**C**) Condensin with an Smc2−Brn1 fusion was incubated with nicked circular DNA as in (A) and the DNA retained after washing with 0.5 M NaCl quantified in relation to unmodified condensin +ATP (mean ± s.d., *n* = 4). (**D**) Condensin with an Brn1−Smc4 fusion as in (C). (**E**) Unmodified condensin or condensin with a cysteine pair for hinge cross-linking (Smc2_K609C_; Smc4_V721C_) were incubated with dibromobimane (+bBBr) or DMSO solvent prior to addition of nicked circular DNA in the presence of ATP. The amounts of DNA retained after a 0.5 M NaCl wash were quantified as in (C) (mean ± s.d., *n* = 3).

Entrapment of DNA in a confined space is a widespread strategy to achieve processivity of enzymes with dynamic nucleic acid interactions, including DNA polymerase sliding clamps or replicative helicases (*19*), damage repair enzymes MutS (*20*) and Rad50 (*21*), type-II topoisomerases (*22*) or the bacterial motor protein FtsK (*23*). Biochemical and structural evidence support the notion that cohesin (*15-17, 24-26*) and condensin (*27*) topologically constrain DNA, but thus far fell short in revealing whether, and if so how, DNA entrapment can form, let alone enlarge, DNA loops. Here, we reconstituted the loading of active condensin complexes onto DNA, which enabled us to reconstruct their reaction cycle at molecular detail. We identified chambers within the protein complex that encircle the static and translocating segments of a growing DNA loop and resolved their DNA interactions at near-atomic resolution. Remarkably, disruption of the bicameral separation turns condensin from a strictly unidirectional into a bidirectional DNA loop-extruder. Based on these data, we propose a ‘hold-and-feed’ reaction cycle that explains directional DNA loop extrusion by SMC protein complexes.

### Condensin loads topologically onto DNA without SMC-kleisin ring opening

To define how the condensin complex binds DNA, we developed an *in vitro* system that recapitulates the salt-resistant topological interaction of condensin-chromatin complexes isolated from cells (*27*). We incubated purified *Saccharomyces cerevisiae* (*Sc*) holo condensin with circular plasmid DNA in the presence of ATP and isolated the resulting complexes by immunoprecipitation (**Fig. 1A**). A subsequent high-salt wash (0.5 M NaCl) eliminated linear DNA (fig. S1A), which by its nature cannot be topologically confined. Only circular DNA molecules bound in a salt-resistant manner (**Fig. 1B**) and their formation strictly depended on ATP binding and hydrolysis by condensin (fig. S1B). Whereas relaxation of super-helical tension in circular DNA by nicking one strand of the double helix did not affect salt-resistant binding, linearization by endonuclease (*Xho*I) cleavage just prior to or during high-salt washes efficiently released DNA (**Fig. 1B**). We conclude that the interaction between DNA and condensin in the salt-resistant complexes reconstituted from purified components is topological in nature.

The lumen of the Smc2–Smc4–Brn1^kleisin^ ring creates a self-contained space (SMC-kleisin) that seems ideally suited to topologically entrap DNA, which might enter this space upon ATP-dependent dissociation of the Smc2–Brn1 interface (*18*). However, condensin complexes with a covalent peptide fusion of Smc2 to Brn1 (fig. S2A) still formed salt-resistant complexes with circular DNA (**Fig. 1C**), extruded DNA loops with similar efficiency (123/167 DNAs) and rates as their non-fused counterparts (103/127 DNAs) (fig. S3A, movie S1) and supported cell proliferation in *S. cerevisiae* (fig. S4). Similarly, peptide linker fusion of Brn1 to Smc4 (fig. S2B) neither abolished the *in vitro* formation of salt-resistant condensin-DNA complexes (**Fig. 1D**) nor affected DNA loop extrusion efficiencies (176/246 DNAs analyzed) or rates (fig. S3B, movie S1) and supported condensin *in vivo* function (fig. S4). Dibromobimane (bBBr) cross-linking of cysteine residues engineered into the Smc2–Smc4 hinge domains (fig. S2C) did also not impair the formation of salt-resistant DNA complexes (**Fig. 1E**). Titration experiments with mixtures of wild-type and inactive mutant (QL) condensin complexes ruled out that the remaining non-cross-linked complexes were responsible for retaining these DNA molecules (fig. S2D). Together, these results argue against the notion that any of the three subunit interfaces function as a DNA entry gate and call into question whether DNA is at all topologically encircled by the SMC-kleisin ring.

We therefore probed DNA entrapment in the SMC-kleisin ring by analyzing native complexes between condensin and circular mini-chromosomes isolated from yeast cells. We covalently circularized the SMC-kleisin ring by combining the Smc2–Brn1 fusion with cysteine cross-linking the Smc2–Smc4 and Smc4–Brn1 interfaces (fig. S5). Addition of bBBr simultaneously cross-linked both cysteine pairs in ∼20 % of condensin molecules. Yet, unlike for cohesin (*25*), we failed to detect sodium dodecyl sulfate (SDS)-resistant catenanes between the covalently circularized condensin rings and circular mini-chromosomes. Although condensin binds DNA topologically, this topological interaction cannot be explained by a single passage of DNA through the SMC-kleisin ring.

### DNA is pseudo-topologically entrapped in two kleisin chambers

Mapping the connectivity of Brn1^kleisin^ segments in structural models of the ATP-free apo state condensin (*28*) indicated the presence of three alternative chambers, each suited to accommodate a DNA double helix (**Fig. 2A**). Chamber I is created by the first ∼200 residues of Brn1, which bind the Ycs4^HEAT-I^ subunit and contact the Smc2_head_ region. Chamber II is created by a ‘safety belt’ loop of ∼130 Brn1 residues that forms within the groove of the Ycg1^HEAT-II^ solenoid and has already been shown to entrap DNA (*12*). An intermediate (IA) chamber is created by Brn1 stretches that connect Ycs4 to Ycg1 and Ycg1 to Smc4_head_, respectively. Note that all three kleisin chambers are within the SMC-kleisin tripartite ring circumference and are separated by impermanent protein interfaces: Dissociation of Ycs4 from Smc4_head_ (*18*) fuses chambers I and IA, while disengagement of the ‘latch’ and ‘buckle’ segments of the Brn1 safety belt (*12*) fuses chambers IA and II.

**Fig. 2.**
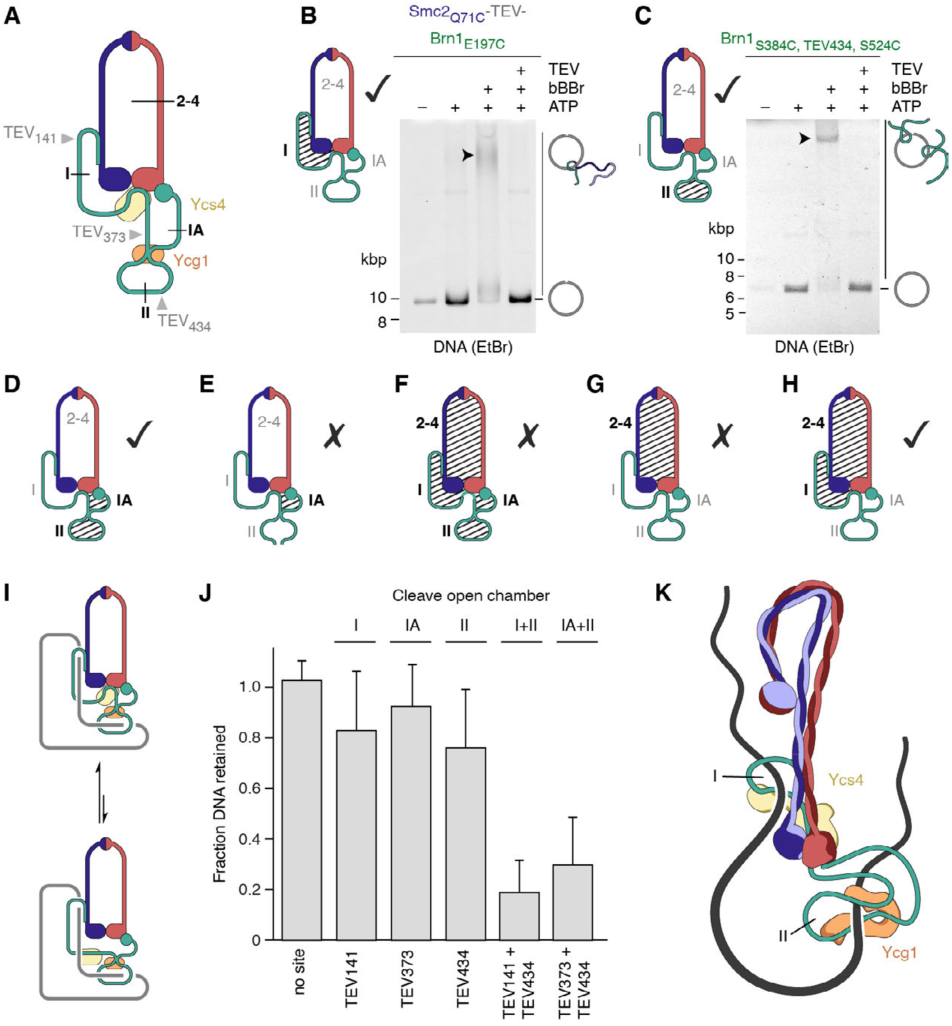
Condensin constrains DNA in two kleisin chambers. (**A**) Schematic representation of condensin in the ATP-free state. Kleisin chambers I, IA and II and the positions of engineered TEV target sites are indicated. (**B**) Covalent circularization of chamber I (shaded area) by cysteine cross-linking (Smc2_Q71C_; Brn1_E197C_) of the Smc2−Brn1 fusion protein. Agarose gel electrophoresis mobility shift of SDS-resistant condensin-DNA catenanes revealed by ethidium bromide staining. (**C**) Electrophoresis as in (B) after covalent circularization of chamber II (shaded area) by cysteine cross-linking (Brn1_S384C, S524C_). (**D–H**) Additional configurations tested for the formation of SDS-resistant catenanes. Checks and crosses indicate whether catenanes were detected or absent, respectively (fig. S6). (**I**) Schematic configuration of the condensin-DNA complex with equilibrium of chamber I and IA fusion. (**J**) Quantitation of salt-resistant condensin-DNA complexes retained after cleavage at indicated TEV sites within Brn1 (mean ±s.d., *n* ≥ 3). (**K**) Model of the DNA path through the ATP-free apo condensin complex.

We systematically explored the involvement in DNA binding of the three Brn1 chambers and the Smc2–Smc4 lumen by covalent closure of single or combinations of multiple chambers using bBBr cross-linking after condensin had been loaded onto circular DNA *in vitro*. Note that these experiments probed the nucleotide-free apo state of the complex, since ATP supplied for the loading reaction was washed away prior to cross-linking. Closure of Brn1 chamber I (fig. S6A), of chamber II (fig. S6B) or of combined chambers IA and II (fig. S6C) produced SDS-resistant DNA–condensin catenanes that were again resolved by opening with TEV protease cleavage (**Fig. 2B–D**). Similar strategies to circularize chamber IA alone (fig. S6D), the entire Smc2–Smc4– Brn1 ring (fig. S6E) or the Smc2–Smc4 lumen (fig. S6F) failed to produce SDS-resistant catenanes (**Fig. 2E–G**), in contrast to a combination that created a circularized compartment between the Smc2–Smc4 lumen and kleisin chamber I (**Fig. 2H**, fig. S6G).

The only configuration that meets the restraints set by these results (fig. S7) places a DNA loop enclosed simultaneously by chambers I and II into the apo conformation of the complex (**Fig. 2I**). We confirmed that DNA is entrapped in both Brn1 chambers at the same time by opening chambers either individually or in combination with site-specific TEV cleavage (fig. S7A). While opening individual chambers had only minor effects, opening of chambers I or IA in combination with chamber II released the majority of bound DNA (**Fig. 2J**, fig. S8B). Note that the low affinity (K_d_ = 0.63 µM (*18*)) of the Ycs4–Smc4_head_ interaction that separates chambers I and IA will allow escape of DNA entrapped in chamber I through a gap created in chamber IA during the extended incubation period required for TEV protease cleavage (**Fig. 2I**). The ‘pseudo-topological’ entrapment of a DNA loop in the SMC-kleisin ring as depicted in **Fig. 2K** explains why none of its interfaces needs to open for DNA entrapment and why ring circularization does not produce denaturation-resistant DNA catenanes – in contrast to cohesin involved in sister chromatid cohesion, which encircles DNA in a truly topological manner (*24, 25*).

### Cryo-EM of ATP-bound condensin reveals the role of DNA in kleisin chambers

To gain detailed insight into the fate of bound DNA in kleisin chambers I and II after ATP binding, we trapped a hydrolysis-deficient version (EQ) of the *Sc* condensin holo complex in presence of 50-bp DNA duplexes and determined its structure by cryo-EM. Single-particle analysis revealed a high degree of flexibility among individual molecules. Neural network-based particle picking combined with 3D classification procedures identified two well-ordered yet flexibly linked modules, each bound to a DNA duplex (fig. S9, S10). The quality of cryo-EM reconstructions of each module allowed *de novo* model building for both modules (fig. S11, table S1), facilitated by high-resolution crystal structures of the individual condensin subunits (*12, 18*).

The catalytic ‘core’ module is composed of Smc2_head_ and Smc4_head_ domains bound to the Ycs4^HEAT-I^ subunit (**Fig. 3A**), whereas the ‘peripheral’ module contains the Ycg1^HEAT-II^ subunit (**Fig. 3B**). Our cryo-EM reconstructions furthermore allowed unambiguous tracing of the Brn1^kleisin^ through the entire complex: Ordered segments of Brn1 ranging from its amino-terminal helix-turn-helix domain (Brn1_N_) to its carboxy-terminal winged helix domain (Brn1_C_) thread through both modules. Disordered linker regions connect the segments and consequently flexibly tether the two modules in the DNA-bound state. At both DNA binding sites, the only conceivable paths of the linker regions lead over the bound double helices. Thus, our findings provide a structural basis for understanding the key role of the kleisin subunit for condensin function: Brn1 mediates strategic inter-subunit interactions throughout the complex and simultaneously establishes the formation of two separate, yet flexibly linked chambers that topologically entrap DNA.

**Fig. 3.**
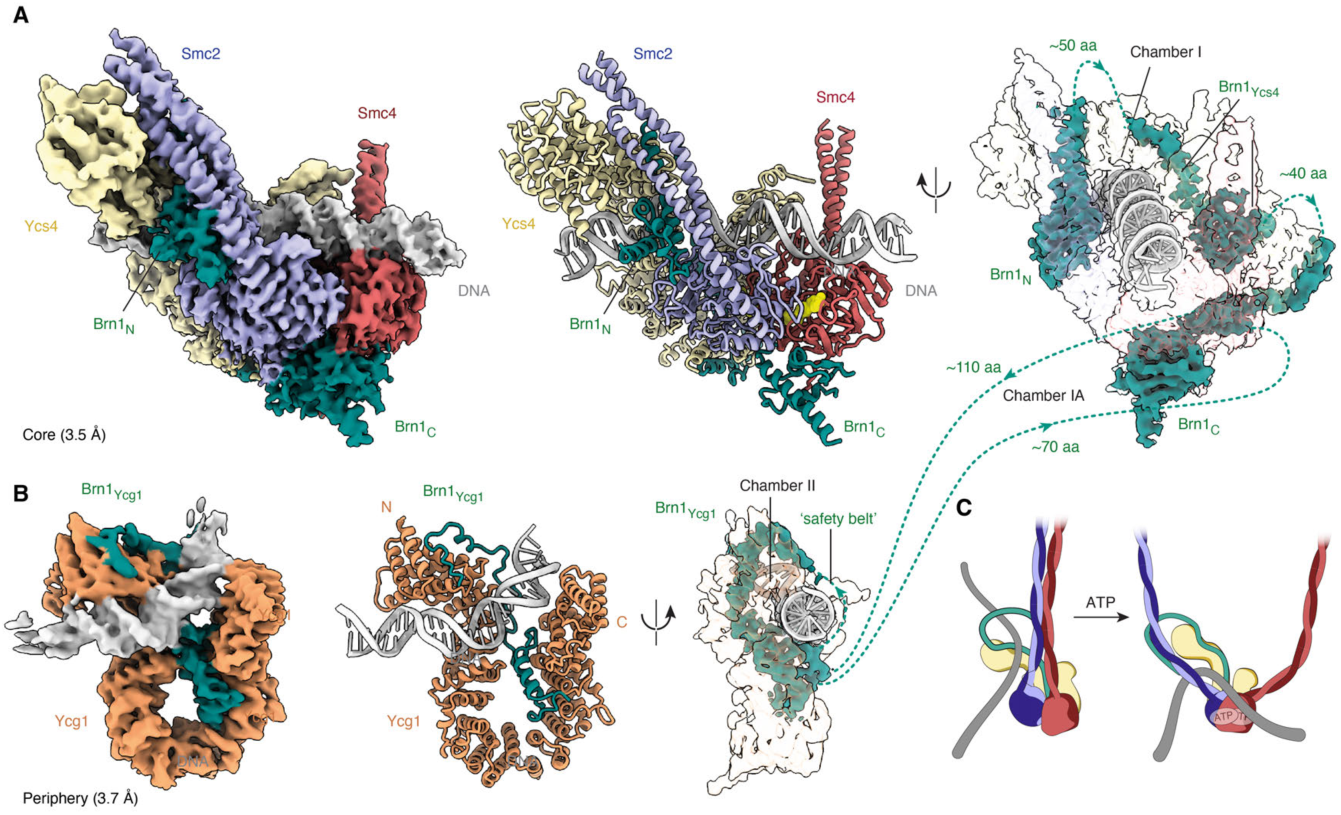
Cryo-EM structure of ATP-engaged condensin with DNA bound in both kleisin chambers. (**A**) Density map and models of the DNA-bound condensin core subcomplex composed of Smc2_head_ (blue), Smc4_head_ (red), Ycs4 (yellow) and Brn1 (green) resolved to a nominal resolution of 3.5 Å. (**B**) Density maps and models of the DNA-bound condensin peripheral subcomplex composed of Ycg1 (orange) and Brn1 (green) resolved to a nominal resolution of 3.7 Å. Unresolved Brn1 connections are indicated as dotted lines. (**C**) Schematic representation of the tilting motion of DNA entrapped in kleisin chamber I upon ATP binding.

A comparison to nucleotide-free apo condensin (*28*) identifies profound conformational rearrangements at the core module, which forms chamber I. Engagement of Smc2_head_ and Smc4_head_ domains by sandwiching ATP at both active sites (fig. S12A) results in a swivel motion, which increases the opening angle between the coiled coils by ∼25° to create an open V shape (fig. S12B), resulting in a highly dynamic, opened lumen between the unzipped coils. Ycs4 undergoes a large conformational change (fig. S13), most likely caused by multivalent interactions with Brn1_N_, the Smc2 coiled coil and approximately half of the 38 visible base pairs of DNA that are accommodated in the positively charged groove on the concave side of its HEAT-repeat solenoid (fig. S14A). We confirmed the importance of these DNA interactions for *in vivo* condensin function (fig. S14C), DNA-dependent ATPase stimulation (fig. S14D) and DNA loop extrusion (fig. S14E). Homologous DNA interactions are also conserved for cohesin (*15-17*). Although the ATP-free apo structure of condensin adapts a markedly different conformation, most of the local surface of Ycs4 that contacts the DNA backbone remains accessible and unchanged in the absence of nucleotide (fig. S14B), supporting the conclusion that kleisin chamber I also entraps DNA in the ATP-free state.

Taken together, our cryo-EM structures in conjunction with biochemical mapping reveal that the concerted opening the of coiled coils from a tightly zipped (*28*) into an open configuration together with a clamping motion of Ycs4 presumably pushes the DNA in chamber I onto the newly formed binding surface of the engaged Smc2_head_ and Smc4_head_ domains (**Fig. 3C**, fig. S15). This previously unanticipated power-stroke movement elegantly explains how ATP binding might fuel the motor function of condensin by feeding a new DNA loop segment into the inter-coil lumen (see below).

The peripheral module visualizes the structure of kleisin chamber II, which is created by Ycg1 bound to the Brn1 safety-belt segment and flexibly linked to the catalytic ‘core’. Whereas a comparison with previous crystal structures shows no major conformational rearrangements of the protein subunits (*12, 29*), the DNA double helix sharply bends almost 90° as it binds to a newly formed composite interface formed by Brn1 and the Ycg1 HEAT-repeat solenoid (fig. S16). This deformation might provide chamber II with the ability to resist longitudinal pulling forces acting on the bound DNA, consistent with a possible anchoring function.

### The kleisin chambers provide anchor and motor functions for DNA loop extrusion

Asymmetric DNA loop extrusion by condensin requires that a single complex must grasp both, the immobile (‘anchor’) and translocating (‘motor’) DNA segments at the stem of the expanding loop (*6*). If the two identified kleisin chambers were – at least during part of the reaction cycle – responsible for these two functions, release of DNA from the motor chamber should retain condensin on the DNA position where extrusion was initiated. Release of DNA from the anchor chamber should, in contrast, retain condensin at the motor end of the original loop, distal from where loop extrusion started.

We followed the fate of condensin complexes labeled with an ATTO647N fluorophore in single-molecule DNA loop extrusion assays in the presence of TEV protease (**Fig. 4A**). Non-cleavable condensin on DNA loops that ruptured spontaneously was, in most cases (55/59 dissolved loops), retained where loop extrusion had originated and in the remaining few cases (4/59) dissociated upon loop rupture (**Fig. 4B**, fig. S17A, movie S3). We confirmed that condensin remained anchored at its starting position when loops snapped on DNA molecules arched by side-flow (fig. S17B). Spontaneous liberation of condensin-mediated DNA loops thus primarily involves release of DNA from the motor entity, occasionally from both motor and anchor, but never from the anchor entity alone.

**Fig. 4.**
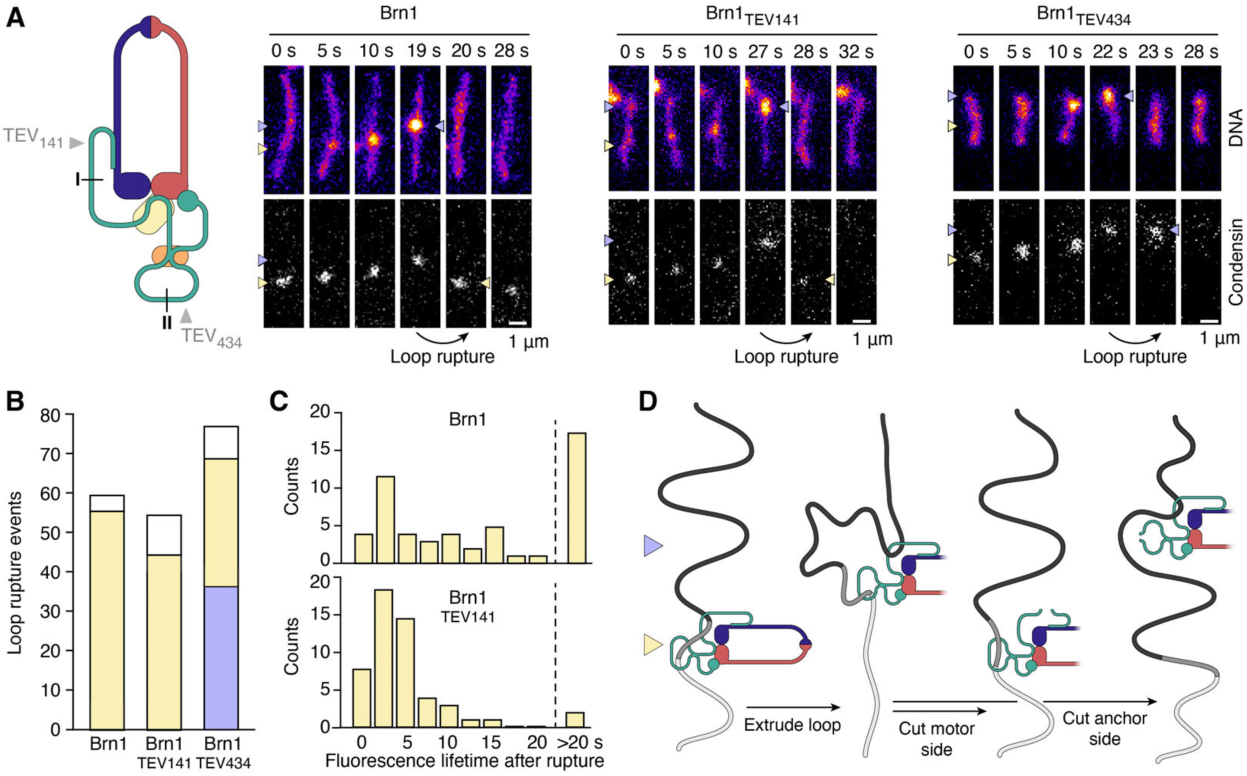
Identification of motor and anchor chambers. (**A**) Single-molecule DNA loop extrusion on SxO-stained surface-tethered λ-phage DNA (48.5 kbp) molecules by ATTO647N-labeled condensin with TEV cleavage sites introduced in chambers I or II of Brn1. Starting and end positions of the DNA loop are highlighted with yellow and blue arrowheads, respectively. Scale bar = 1 µm. (**B**) The position of condensin 0.5–1 s after DNA loop rupture was scored as non-detectible (white), back at the loop start site (yellow) or on the translocating end of the loop (blue). (**C**) Histogram of ATTO647N-condensin fluorescence lifetimes at the loop start site after loop rupture. (**D**) Schematic representation of experiment and results.

DNA loops created by condensin with a TEV cleavage site in chamber I released in a similar manner as spontaneous rupture events (**Fig. 4B**, fig. S17A, movie S4), with condensin retained at the anchor position (44/54) or lost from the DNA (10/54). These events were attributable to opening of chamber I, since we detected the ATTO647N fluorophore attached to Brn1_N_, which is released from the complex upon TEV cleavage *(18)*, for a considerably shorter time than after spontaneous rupture of non-cleavable condensin (**Fig. 4C**). We conclude that opening of kleisin chamber I releases the motor segment of the DNA loop.

In stark contrast, when loops made by condensin with a TEV cleavage site in kleisin chamber II dissolved, condensin was released from the anchoring position and retained at the translocating site in nearly half of the observed cases (36/76; **Fig. 4B**, fig. S17A, movie S5). In rare instances, condensin continued to translocate in the same direction after loop rupture, now trailing a small DNA density it was no longer able to expand (fig. S17C). We observed several cases of condensin translocation without DNA loop expansion after prolonged incubation with TEV protease (fig. S17D). Consistent with the previous finding that mutation of the kleisin safety belt results in DNA loop slippage *(6)*, our experiments demonstrate that chamber II creates the anchor segment of the DNA loop. The remaining loop rupture events, where condensin remained at the anchor position (32/76) or dissociated (8/76), presumably correspond to spontaneous loop ruptures, which we still expect to occur with TEV-cleavable condensin.

### The anchor chamber defines DNA loop extrusion directionality

Our TEV cleavage experiments imply that anchor and motor activities of condensin can be functionally separated. We were able to generate a separation-of-function version for the condensin complex from the filamentous fungus *Chaetomium thermophilum* (*Ct*) (fig. S18A), which displays DNA-stimulated ATPase activity at temperatures up to 50 °C (fig. S18B) and retains much of its affinity for DNA even in the absence of Ycg1, in contrast to *Sc* condensin (fig. S18C).

*Ct* holo condensin induced local DNA compaction events on tethered DNA molecules (**Fig. 5A**) that emerged as DNA loops upon changing the direction of buffer flow (fig. S19A). DNA loop formation required ATP and Mg^2+^ and was abolished by mutation of the Smc2 and Smc4 ATP-binding sites (QL) (fig. S19B). Remarkably, *Ct* ΔYcg1 condensin initiated the formation DNA loops (**Fig. 5B**) with even greater efficiency than *Ct* holo condensin (**Fig. 5C**). Only when we in addition deleted (Brn1_Δ515–634_) or mutated conserved positively charged residues within the Brn1 safety belt loop (Brn1_BC_) did we no longer observe loop extrusion. Quantitation of the DNA loop extrusion parameters revealed that *Ct* ΔYcg1 condensin generated loops at similar rates (**Fig. 5D**). Yet, the lifetime of loops generated by *Ct* ΔYcg1 condensin was significantly increased when compared to loops generated by *Ct* holo condensin (**Fig. 5E**), which otherwise snapped soon after the complex reached the stall force for loop extrusion (**Fig. 5F**). We conclude that kleisin chamber II, but not the presence of Ycg1, is essential for condensin-mediated DNA loop extrusion.

**Fig. 5.**
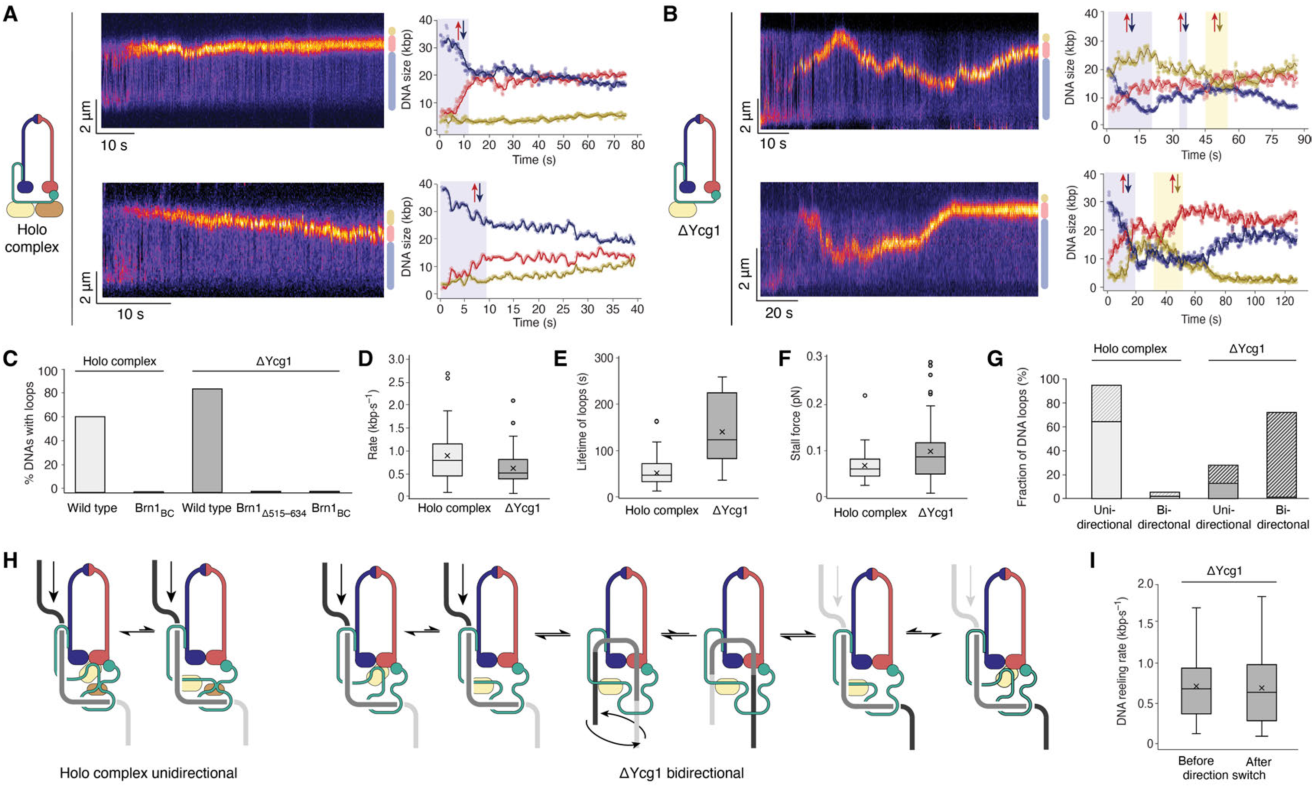
Merge of DNA chambers enables condensin-mediated DNA loop extrusion to change direction. (**A**) Sample kymographs of DNA loop extrusion by *Ct* holo condensin on λ-phage DNA stained with Sytox orange. Fluorescence intensity plots represent DNA fluorescence above (yellow) or below (blue) the extruded loop (red). (**B**) Sample kymographs of DNA loop extrusion by *Ct* ΔYcg1 condensin as in (A). (**C**) Fraction of DNA molecules displaying loops created by *Ct* holo (wild type or Brn1_BC_) and ΔYcg1 (wild type, Brn1_Δ515–634_ or Brn1_BC_) condensin (*n* = 509, 158, 307, 70 or 145 DNAs analyzed). (**D**) Box plot of DNA loop extrusion rates. Lines indicate median, crosses indicate mean, boxes indicate first and third quartile, whiskers mark the median ± 1.5 (third quartile – first quartile). (**E**) Box plot of lifetime of DNA loops as in (D). (**F**) Box plot of stall forces as in (E). (**G**) Fraction of unidirectional or bidirectional DNA loop extrusion events observed for *Ct* holo and ΔYcg1 condensin (*n* = 56, 79). Shaded areas indicate events that displayed anchor slippage. (**H**) Illustration of a strict separation of motor and anchor DNA segments in holo condensin (left) or exchange of segments in the absence of Ycg1 (right). (**I**) Loop extrusion rates before and after direction switch by *Ct* ΔYcg1 condensin (mean ± s.d., *n* = 56).

Like condensin from other species (*6, 9*), *Ct* holo condensin almost exclusively reeled in DNA unidirectionally (53/56 DNA loops; **Fig. 5G**, fig. S20A, movie S6). In contrast, *Ct* ΔYcg1 condensin frequently switched directions during loop extrusion (57/79; **Fig. 5G**, fig. S20B, movie S7). On some DNA molecules, the DNA loop changed direction as many as six times within a 120-seconds imaging window (fig. S20B, movie S8). The changes in direction were sometimes difficult to discern when they overlapped with anchor slippage events, which were more frequent for DNA loops generated by *Ct* ΔYcg1 condensin than for holo condensin (**Fig. 5G**), but could clearly be identified in the majority of cases when the loop size further increased as condensin reeled in DNA from the opposite direction (fig. S21A). The change in loop extrusion direction is hence not simple backtracking of condensin’s motor entity. It can also not be explained by the action of a second condensin complex that moves into the opposite direction, since such an event would have resulted in the formation of Z-loop structures, which are easily recognizable by the elongated DNA density (*30*) and were rare under the conditions of our assay (fig. S21B).

We propose that the observed turns instead reflect an exchange of motor and anchor DNA segments within the extruding condensin complex (**Fig. 5H**). If this were the case, the speed of loop extrusion should be identical in either direction. Loop extrusion rates after switching direction were indeed very similar to the original translocation rates (**Fig. 5I**).

### A hold-and-feed mechanism drives SMC-mediated DNA loop extrusion

SMC complexes stand out from conventional DNA motor proteins by their ability to translocate in steps of kilo-base pairs in length (*6-8*). Current models for the molecular mechanism of DNA loop extrusion fail to explain how consecutive steps can proceed in a directional manner on a DNA substrate that lacks intrinsic polarity (*31*). Biochemical mapping of the path of DNA through two kleisin chambers (**Fig. 2**), structures of the identical protein complex in ATP-free *(28)* and ATP-bound (**Fig. 3**) states and the assignment of motor and anchor functions to the DNA binding sites by single-molecule imaging (**Fig. 4**) now provide the foundation for a mechanistic description of the SMC-mediated DNA loop extrusion cycle:

The DNA-segment-capture model (*32*) proposes that SMC dimers grasp DNA loops generated by random thermal motion. Our data instead suggest that the concerted tilting of a DNA double helix that is entrapped in kleisin chamber I actively feeds DNA in-between the unzipped coiled coils upon ATP-mediated SMC head engagement (**Fig. 6A**, fig. S22A). As a result of this power-stroke motion, two DNA loops are now pseudo-topologically entrapped by the condensin complex (**Fig. 6B**). To reset the complex to the apo state following nucleotide hydrolysis, SMC head disengagement most likely first results in the ‘bridged’ conformation observed previously (*28*). As a consequence, the head-proximal segment of the newly captured loop releases from kleisin chamber I (**Fig. 6C**). Simultaneously, zipping up of the SMC coiled coils (*32, 33*) and/or tilting of the latter onto the folded coils (*28, 34*) move the distal loop segment towards the ATPase heads, where it remains confined between HEAT-repeat subunit I and the SMC coiled coils. To regenerate the initial conformation with DNA in kleisin chamber I, this DNA segment then merely has to tilt into the DNA-binding groove of the HEAT-repeat subunit, which is only possible in one direction due to geometric restrictions.

**Fig. 6.**
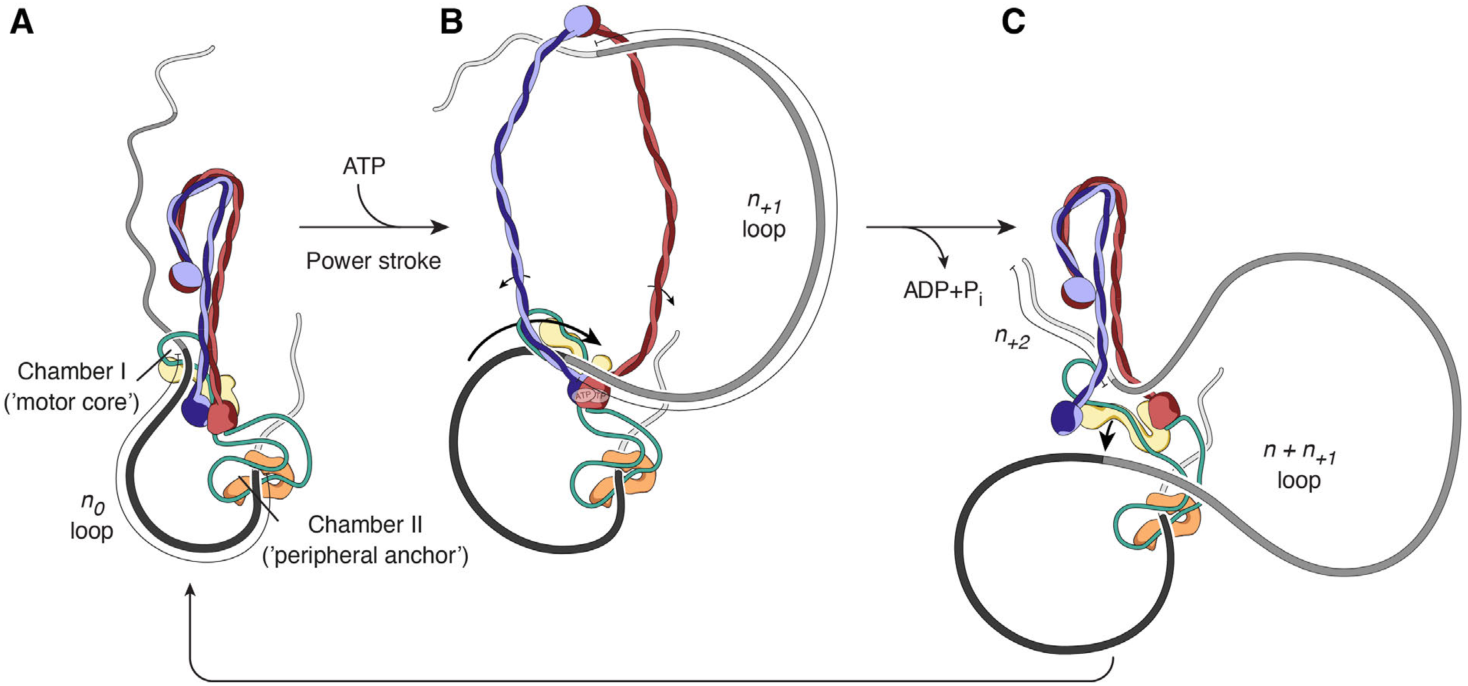
A hold-and-feed mechanism for SMC-mediated DNA loop extrusion. (**A**) After loading, ATP-free condensin entraps a DNA loop (*n*_*0*_) pseudo-topologically between kleisin chambers I and II. (**B**) ATP binding drives SMC head dimerization, coiled-coil opening and Ycs4^HEAT-I^ repositioning to feed DNA held in kleisin chamber I in-between the SMC coiled coils at the SMC motor core (power stroke). The result is the pseudo-topological entrapment of a new DNA loop (*n*_*+1*_) in the coiled-coil lumen. (**C**) ATP hydrolysis drives the transition into the ATP-free ‘bridged’ state of the SMC motor to release one *n*_*+1*_ DNA segment, while the peripheral anchor remains bound upstream. This step merges both loops (*n*_*0*_ *+ n*_*+1*_). Return to the ground state configuration repositions the remaining *n*_*+1*_ DNA segment into chamber I. Condensin is now ready to extrude the next DNA loop (*n*_*+2*_). Fig. S22 summarizes structural information supporting each step as well as possible intermediates of the reaction cycle.

DNA entrapment in kleisin chamber I hence ensures that translocation proceeds processively and in a single direction, always threading the next DNA segment into the SMC coiled-coil lumen from the same end of the DNA loop. This translocation mechanism explains how condensin can step over flexibly tethered obstacles that are many times its size (*35*), while nevertheless binding DNA pseudo-topologically (fig. S22B).

DNA entrapment in kleisin chamber II is essential for anchoring condensin to DNA (*6, 12*), whereas HEAT-repeat subunit II is not (**Fig. 5**). This finding is inconsistent with a recently proposed Brownian ratchet mechanism as the fundamental principle for DNA loop extrusion by SMC complexes (*36*). The HEAT-repeat subunit is, however, required to close off the kleisin safety belt and thereby separate anchor and motor strands of the DNA loop, since its absence from the condensin complex turns an exclusively unidirectional DNA loop extruder into one that frequently switches direction. The natural merge of chambers II and IA in the absence of a kleisin safety belt in cohesin (*12, 13*) presumably allows for a frequent exchange of motor and anchor strands (*7, 8*), which explains how monomeric cohesin can extrude DNA loops bidirectionally. Binding of the cohesin HEAT-II subunit to the CTCF boundary factor most likely prevents strand exchange and thereby provides a molecular account for the CTCF convergence rule for topologically-associating domains (*37*). Confinement of the DNA in two kleisin chambers thus not only forms the basis of DNA translocation but also dictates the directionality of loop extrusion by SMC protein complexes.

## Acknowledgments

We thank Jenny Ormanns and Jutta Metz for assistance with the generation of yeast strains, Shveta Bisht for protein purification, Fabian Merkel for advice and help with cryo-EM, Robin Stipp for help setting up insect cell expression, Thomas Hoffmann for scientific computing support, Marko Lampe of the EMBL Advanced Light Microscopy Facility, Felix Weis of the EMBL Cryo-Electron Microscopy Platform and the Proteomics Core Facilities.

## Author contributions

IAS, SD, MK and SB purified condensin complexes; IAS performed *in vitro* DNA loading assays; IAS and SD performed single-molecule experiments; LL and SE performed cryo-EM experiments and processed data; LL, SE and MH built structure models; IAS and CS performed yeast experiments; SE and CHH supervised the work and acquired funding; IAS, SE and CHH wrote the manuscript with input from all authors.

## Data and materials availability

All data are available in the main text or the supplementary materials. Plasmids, yeast strains and image analysis scripts will be made available upon request. Coordinates of DNA-bound condensin are available in pdb under accession numbers xxx (core) and xxx (periphery).

## Supplementary Materials

### Materials and Methods

#### *Purification of* Saccharomyces cerevisiae (Sc) *condensin*

Wild-type and mutant *Sc* condensin complexes were essentially expressed and purified as described previously (*38*). All five subunits were expressed from two 2µ-based high copy plasmids under the control of galactose-inducible promoters transformed into *S. cerevisiae*. One plasmid contained *pGAL10-YCS4* (or mutations thereof) *pGAL1-YCG1 TRP1*, and the other *pGAL7-SMC4-StrepII*_*3*_ *pGAL1-SMC2 pGAL1-BRN1-His*_*12*_*-HA*_*3*_ *URA3* or their derivates, including fusion proteins (see table S2). Cultures were maintained at 30 ºC in –Trp–Ura media and 2 % (w/v) D-glucose and overexpression was induced by transfer to –Trp–Ura media and 2% (w/v) raffinose for 6 h and addition of 2 % (w/v) D-galactose for 16 h. Cell lysates were prepared in a FreezerMill (Spex) in buffer A (50 mM Tris-HCl pH 7.5, 200 mM NaCl, 5 % (v/v) glycerol, 5 mM β-mercaptoethanol, 20 mM imidazole) supplemented with cOmplete EDTA-free protease inhibitor mix (11873580001, Merck). Lysates were cleared by centrifugation, loaded onto a 5-mL HisTrap FF column (Cytiva) and eluted with 220 mM imidazole in buffer A. The eluate fractions were supplemented with 0.01 % (v/v) Tween-20, 1 mM EDTA and 0.2 mM PMSF and incubated ∼16 h with Strep-Tactin Superflow high capacity resin (2-1208-010, IBA), eluted with buffer B (50 mM Tris-HCl pH 7.5, 200 mM NaCl, 5 % (v/v) glycerol, 1 mM dithiothreitol (DTT)) containing 10 mM desthiobiotin (D1411, Merck). For the Brn1–Smc4 fusion condensin, this step was omitted. The eluate was concentrated by ultracentrifugation before size-exclusion chromatography on a Superose 6 increase 10/300 column (Cytiva) pre-equilibrated in buffer B supplemented with 1 mM MgCl_2_. Purified protein was snap-frozen and stored at –80 ºC until use.

#### *Purification of* Chaetomium thermophilum (Ct) *condensin*

*Ct* condensin holo complex and ΔYcg1 tetrameric complex were expressed in Sf21 cells using the Multibac expression system (*39*). Sf21 cells were maintained in SF 900 III serum-free medium (12658019, ThermoFisher) at 27 ºC. Cell pellets were resuspended in buffer A supplemented with cOmplete EDTA-free protease inhibitor mix and sonicated at 4 ºC for 5 cycles of 45 s each in pulse mode, 1 s on, 1 s off, at 50 % energy (SFX-550, Branson). The lysate was cleared by centrifugation at 45,000 × *g* for 1 h and then incubated with Ni Sepharose (175268, Cytiva) beads at 4 ºC for 3 h. After washing the beads with 40–50 column volumes (cv) buffer A (50 mM Tris-HCl pH 7.5, 200 mM NaCl, 5 % (v/v) glycerol, 20 mM Imidazole, 1 mM DTT), proteins were eluted in buffer A containing 300 mM imidazole. The buffer of the eluate fractions was exchanged to low salt buffer (25 mM Tris-HCl pH 7.5, 100 mM NaCl, 5 % (v/v) glycerol, 1 mM DTT) using a desalting column before loading onto a RESOURCE Q (Cytiva) anion exchange column pre-equilibrated with low salt buffer. After washing with 3 cv low salt buffer, proteins were eluted by increasing NaCl concentrations to 1 M in a linear gradient of 60 mL. Peak fractions were pooled and concentrated by ultrafiltration (Vivaspin 30,000 MWCO, Sartorius) before loading onto a Superose 6 increase 10/300 column (Cytiva) pre-equilibrated in buffer B. Peak elution fractions were concentrated by ultrafiltration, snap-frozen and stored at –80 ºC until use. Purified proteins were analyzed by SDS-PAGE and the stoichiometry of the complex was determined by mass spectrometry.

#### Fluorescent labeling of purified condensin complexes

Coenzyme A-coupled dyes were generated as described (*6*) by incubation of a 1.1-fold molar excess maleimide dye-conjugate ATTO647N-maleimide (AD 647N-41, ATTO-TEC) or Alexa Fluor 488 C_5_ maleimide (A10254, ThermoFisher) with Coenzyme A (C3144, Merck) in 1:1 DMSO in 100 mM sodium phosphate pH 7.0 (aq; degassed) buffer at room temperature. A 2-fold molar excess of tris(2-carboxyethyl)phosphine (646547, Sigma) was added to the reaction after 15 min and the reaction was quenched after one hour by addition of a 7-fold molar excess DTT.

Enzymatic site-specific covalent labeling of the Serine hydroxyl of ybbR peptide tags (GTD<u>S</u>LEFIASKLA) on condensin holocomplexes (2-7 µM) was performed in buffer B supplemented with 10 mM MgCl_2_, 1.2 µM Sfp phosphopantetheinyl transferase (P9302, New England Biolabs) and a 5-fold molar excess CoA-dye conjugate reaction mix. After 16 h incubation at 6 ºC, complexes were purified by size-exclusion chromatography on a Superose 6 increase 3.2/300 column (Cytiva) pre-equilibrated in buffer B containing 1 mM MgCl_2_. Peak elution fractions were concentrated by ultrafiltration, snap-frozen and stored at –80 ºC until use.

#### In vitro *DNA loading assays*

DNA substrates were generated from a 6.4-kb plasmid containing *Sc CEN4 TRP1 ARS1 RDN37* sequences and 21 *tetO* repeats (*27*) by incubation with Nb.*Bbv*CI (R0631, New England Biolabs) or *Eco*RI (R3101, New England Biolabs), phenol:chloroform extraction and ethanol precipitation. Loading reactions (30 µL volume) were assembled on ice with 0.1 µM condensin, 250 or 500 ng of substrate DNA and 0.1 g·L^−1^ BSA in reaction buffer (50 mM Tris-HCl pH 7.5, 125 mM NaCl, 50 mM KCl, 5 mM MgCl_2_, 5 % (v/v) glycerol, 1 mM DTT) before addition of 1 mM ATP and 5 min incubation at 25 ºC. Protein-DNA complexes were immunoprecipitated for 20 min on ice with 12.5 µL magnetic protein A Dynabeads (10002D, ThermoFisher) pre-charged with 3 µg 12CA5 anti-HA monoclonal antibody. Beads were washed three times with 100 µL in reaction buffer, before three 0.5 M NaCl washes of 100 µL reaction buffer supplemented with 325 mM NaCl each. Where indicated, the washes were collected and DNA ethanol-precipitated for analysis. Where indicated, DNA was linearized by addition of 1 µL *Xho*I endonuclease (R0146, New England Biolabs) for 20 min at 25 ºC in 30 µL reaction buffer prior to 0.5 M salt washes. In experiments with TEV protease, all samples were incubated for 25 min at 25 ºC either with or without 1.5 µg TEV protease just prior to 0.5 M salt washes. Proteins and DNA-protein complexes were eluted for 10 min at 65 ºC in reaction buffer containing 1 % (w/v) SDS and resolved on 0.75% agarose gels in TAE buffer containing 0.5 mg·L^−1^ ethidium bromide at 1–4 V·cm^−1^ for 16–4 h at 4 ºC. DNA fluorescence was detected on a Typhoon FLA9500 (Cytiva) with a 532-nm laser and 575-nm LP emission filter. All images were analyzed using FIJI (*40*).

#### Dibromobimane (bBBr) cross-linking

In cross-linking experiments, four additional washes of 100 µL each with cross-linking buffer (50 mM sodium phosphate pH 7.5, 125 mM NaCl, 50 mM KCl, 5 mM MgCl_2_, 5 % (v/v) glycerol) were included before cross-linking on ice for 5 min with 100 µL cross-linking buffer containing 0.1 mM TCEP and 0.2 mM bBBr (34025, Sigma) or DMSO. Reactions were quenched with 10 mM DTT. Where indicated, samples were treated with 1.5 µg TEV protease for 25 min at 25 ºC before 0.5 M salt washes. Samples were split 3:1 for agarose gel electrophoresis and SDS-PAGE, respectively. Agarose gels were stained with ethidium bromide after imaging protein fluorescence (Typhoon FLA9500). TEV cleavage efficiencies were determined from SDS-PAGE by in-gel fluorescence detection of the cleaved product in TEV-only controls. Cross-linking efficiencies were determined by in-gel detection of the slower-migrating cross-linked species directly or the reduction in TEV-cleaved product (for Brn1_S384C, TEV434, S524C_).

For cysteine cross-linking at the Smc2–Smc4 hinge (Smc2_K639C_; Smc4_V721C_) prior to DNA incubation, purified protein was first dialyzed against buffer without DTT and then cross-linked with 0.2 mM bBBr (or DMSO) in the presence of 0.1 mM TCEP and reactions quenched with 10 mM DTT.

#### *Mini-chromosome* ex vivo *cross-linking*

Haploid budding yeast strains harboring 2.3-kbp mini-chromosomes (*27*) and expressing circularizable HA-tagged cohesin (C2529) or condensin (C4137) were grown at 25 ºC in –Trp medium to OD_600 nm_ ≈ 1. Cells were collected by centrifugation, washed in PBS at 4 ºC, and resuspended in PBS supplemented with 0.01% (v/v) Triton-X100, 2× cOmplete EDTA-free protease inhibitor mix, 0.1 mM PMSF and 2 mM DTT. Cell lysates were prepared in a FreezerMill (Spex) after snap-freezing the suspension in liquid nitrogen, and cleared by centrifugation (10 + 20) min, 20,000 × *g* at 4 ºC. Complexes were immunoprecipitated with 10 µg anti-HA antibody (12CA5) and 50 µL protein G Dynabeads (10004D, ThermoFisher) per liter culture volume in IP buffer (50 mM Tris-HCl pH 8.0, 100 mM NaCl, 2.5 mM MgCl_2_, 0.25% (v/v) Triton-X100, 1 mM DTT, 1 mM PMSF, 1× cOmplete EDTA-free protease inhibitor mix) for 3 h at 4 ºC. Before cross-linking, beads were washed three times in IP buffer, then three times in cross-linking buffer (25 mM sodium phosphate pH 7.5, 100 mM NaCl, 50 mM KCl, 10 mM MgSO_4_, 0.25% (v/v) Triton-X100). Dibromobimane (0.25 mM) or the equivalent volume of DMSO solvent was added to beads suspended in 100 µL reaction buffer on ice for 10 min and cross-linking was quenched with three washes in cross-linking buffer containing 1 mM DTT. Complexes were eluted for 10 min at 65 ºC in 35 µL elution buffer (50 mM Tris-HCl pH 8.0, 100 mM NaCl, 1 mM MgSO_4_, 1% (w/v) SDS, 1 mM DTT, 1× cOmplete EDTA-free protease inhibitor mix).

For Western Blot, 5 µL of the eluate were supplemented with Laemmli sample buffer and resolved on 3–8% tris-acetate gradient gels (EA0375, ThermoFisher) and transferred onto low fluorescence PVDF (162026, Bio-Rad) by semi-dry blotting (Trans-Blot Turbo, Bio-Rad). HA-tagged subunits were probed with 1 µg anti-HA (12CA5) antibody followed by Cy5-conjugated anti-mouse IgG antibodies (71517515, Jackson Research) and detected on a Typhoon FLA9500 (Cytiva) with a 635-nm laser and 665-nm LP emission filter.

For Southern Blot, the remaining 30 µL of the eluate were supplemented with DNA loading dye (B7024S, New England Biolabs) and DNA was resolved on a 0.75 % (w/v) ultrapure agarose (15895218, ThermoFisher) in TAE buffer in the presence of ethidium bromide (0.5 mg·L^−1^) at 1 V·cm^−1^ and 4 ºC for 24 h, transferred to Hybond-N+ nylon (RPN303B, Cytiva) by capillary force. Mini-chromosomes were hybridized to probes generated with a Prime-It random primer labeling kit (300385, Agilent) and [α-^32^P]-dATP (Hartmann Analytic) on mini-chromosome DNA sequences and detected using a phosphor storage screen on a Typhoon FLA9500 (Cytiva) with a 635-nm laser and 390-nm band pass emission filter.

#### Cryo-EM sample preparation

Purified *Sc* condensin Walker B mutant (Smc4_E1352Q_, Smc2_E1113Q_) holo complexes were diluted to a final concentration of 1 µm in 20 mM HEPES-NaOH pH 7.5, 50 mM NaCl, 5 mM MgCl_2_, 1 mM DTT. ATP was added to a concentration of 1 mM and, after 10 min incubation, 50-bp dsDNA (5’-GTTGACAGTG TCGCAACCTG CACAGGCAAG CTGCTGAGTC TGGTGTAGAC-3’ annealed to the complementary strand) was added to a concentration of 1 µM.

A volume of 3 µl was applied onto a Quantifoil mesh 200 R2/2 grid (Jena Bioscience) after plasma cleaning for 45 s in 25 % oxygen and 75 % argon. Plunge freezing was carried out at 8 °C and 100 % humidity on a FEI Vitrobot Mark IV (ThermoFisher) with a wait time of 2 min, blot force 3 and blot time 1 s.

#### Cryo-EM data collection and processing

Images were recorded on a Titan Krios electron microscope (FEI) equipped with a K2 direct electron detector (Gatan). Images were collected automatically using SerialEM (*41*). A total of 6,544 micrographs were collected in counting mode with total doses of 40 electrons per Å^2^ during exposure times of 8 s, dose fractionated into 40 movie frames at defocus ranges of 1.0–2.2 µm. A magnification of 130,000× resulted in physical pixel size of 1.04 Å.

Cryo-EM data processing was carried out using RELION 3.1.3 (*42*) and CryoSPARC 3.2.0 (*43*). Movie frames were corrected by using 5 × 5 patches and dose weighting in the Motioncor2 implementation of RELION (*44*). CTF parameters were estimated with CTFFIND-4.1 (*45*). Fig. S9 and S10 show an overview of the cryo-EM processing scheme used for analyzing the dataset.

Initial particle picking was done using WARP (version 1.0.9) trained with a BoxNet2Mask model avoiding picking of aggregates and high contrast artifacts (*46*). 2D classification of the entire set of picked particles in CryoSPARC (fig. S9) indicated a large conformational complexity of the condensin-DNA particles. To overcome the challenges for structure determination associated with this complexity, we used a combination of *ab initio* reconstruction and simultaneous 3D classification implemented CryoSPARC, neuronal network-based approaches implemented in Topaz, as we well as 3D classification procedures in RELION.

2D classes were categorized as core 1 (C1), Ycg1/Ycs4 (Y), core 2 (C2) or fragments (F) (fig. S9B). Particles corresponding to C1, Y and C2 were used for generation of *ab initio* reconstructions via simultaneous 3D classification. Notably, *ab initio* reconstructions of C2 particles resembled the previously described nucleotide-free state of condensin (*28*). Particle sets of C1 and Y were used for Topaz neural network training and picking. Neuronal network training was performed in Topaz using 4-fold binned micrographs and based on the conv_127 CNN model (*47*). 3D classification of Topaz-picked particles was performed in RELION using references low-pass-filtered to a resolution 40 Å. Similarly, all 3D references used for 3D template-based picking, 3D classification and 3D refinements were low-pass filtered to 40 Å to avoid high-resolution reference bias.

*Ab initio* classes 1 and 2 of C1 were used as references for 3D classifications on subset of 251,513 particles. Two rounds of 3D classifications were carried out on a subset of 276,846 Y particles using the *ab initio* class 2 as a 3D reference. The 3D classification using the *ab initio* class 2 yielded a cryo-EM density that indicated an alternative, yet highly similar, DNA-binding mode of condensin. However, preferred orientations of the particle projections limited high-resolution reconstruction of particles of this conformation. The resulting C1 and Y densities were 3D-refined (fig. S9F, G) and used for the rest of the analysis.

For C1 particles, the core density (fig. S9F) was used as a 3D template for picking by RELION and led to the extraction of 1,203,041 particles that underwent two consecutive rounds of 3D classifications. An initial 3D refinement was then carried out on a subset of 285,280 particles. The resulting density was used as a reference for a 3D classification using fixed Euler angles. This 3D classification identified 66,205 particles that were used for two rounds of per-particle CTF refinement, Bayesian polishing and final 3D refinement of DNA-bound condensin core complex particles in RELION.

For Y particles, the Ycg1 density (fig. S9G) was used for neural network Topaz picking, using the above-mentioned training parameters and particles. The resulting 571,014 picked particles were 3D classified. 217,204 particles, corresponding to the most populated class, were 3D refined. Finally, one additional 3D classification was performed with fixed Euler angles on the same particle set. This 3D classification identified 114,600 particles that were used for two rounds of per-particle CTF refinement, Bayesian polishing and final 3D refinement of the DNA-bound Ycg1 complex in RELION.

B-factor estimation as well as sharpening was performed according to Rosenthal and Henderson (*48*). Overall resolution estimates were calculated from Fourier shell correlations at 0.143 between the two independently refined half-maps. Local resolution estimation and local resolution filtering of 3D reconstructions was performed using the estimated B-factors as well as procedures implemented in RELION.

#### Structure model building

The model for the core complex was built into a 3.46 Å B-factor-sharpened map. Starting models for the individual subunits were obtained from PDB entries 6YVU and 6YVD (*28*) and rigid body fitted using ChimeraX (*49*). The Ycs4_C_ and the Smc2_neck_–Brn1_N_ regions were fitted individually to accommodate for large conformational changes to the respective starting models. The model was then rebuilt in Coot (*50*) using its interactive sharpening procedures with several parts built *de novo* (e.g. Ycs4_C helix_, Brn1_Ycs4_ and several loop regions). DNA was modelled as a poly-A:T double helix and fitted into the electron density using Geman-McClure restraints in Coot. Regions in the model with poor local resolution (e.g. Ycs4_N_, Brn1_Ycs4_) were aligned with available models from the AlphaFold server (*51*). The model was further improved and finalized by cycles of editing in ISOLDE (*52*) and Coot and automated refinement in phenix.real_space_refine using secondary structure restraints (alpha, beta, base-pair) and Ramachandran restraints (*53*) against a local resolution-filtered, B-factor-sharpened map.

The model for the peripheral complex was built into a 3.69 Å B-factor-sharpened map. The starting model for the peripheral Ycg1 complex was obtained from PDB entry 5OQN (*12*). Ycg1 residues 7–233 and 234–910 and Brn1 were placed individually using rigid body fitting in ChimeraX. The model was rebuilt using interactive sharpening procedures and the DNA was modelled as poly-A:T and fitted into the electron density using Geman-McClure restraints in Coot. Finally, the model was further improved and finalized by cycles of editing in ISOLDE and Coot, and automated refinement in phenix.real_space_refine using secondary structure restraints (alpha, beta, base-pair) and Ramachandran restraints against a local resolution filtered, B-factor sharpened map.

#### ATP hydrolysis assays

Plasmid DNA was relaxed with *E. coli* topoisomerase I (M0301, New England Biolabs) or by nicking with Nb.*Bbv*CI (R0631, New England Biolabs) and purified by phenol:chloroform extraction followed by ethanol precipitation. 10 µL ATP hydrolysis reactions were set up with 0.5 µM *Ct* condensin in the presence or absence of 25 nM relaxed circular 6.4-kb plasmid DNA in ATPase buffer (40 mM Tris-HCl pH 7.5, 125 mM NaCl, 10 % (v/v) glycerol, 5 mM MgCl_2_, 5 mM ATP, 1 mM DTT and 50 nM [α-^32^P]-ATP; Hartmann Analytic). ATP hydrolysis reactions were equilibrated to 25 ºC for 10 min and were initiated by addition of ATP. Upon addition of ATP, the reactions were incubated at different temperatures as indicated (25–50 ºC). 1 µL of the reaction was spotted onto PEI cellulose F TLC plates (Merck) every 2 or 3 min for a total duration of 12 or 18 min. The reaction products were separated by 1 M LiCl and 2 M formic acid solution and the signals were detected via a phosphor-imager screen, which was scanned on a Typhoon FLA 9500 scanner (Cytiva). ATP hydrolysis rate was calculated by linear fit of the ADP/ATP ratio during in the linear range of the reaction. If not stated otherwise, ATP hydrolysis assays for each sample were performed in three replicates.

#### Electrophoretic mobility shift assay (EMSA)

6-FAM-labeled 50-bp dsDNA for EMSA was prepared by annealing 5’-6-FAM-ATTAGTTACT AAGATCCTTC CTCTGTAGAAGAATGAGATTATCGGAGACAG-6-FAM-3’ and the complementary 5’-FAM-labeled oligo, both purified by HPLC (Sigma), at a concentration of 20 µM in a temperature gradient of 0.1 ºC·s^−1^ from 95 ºC to 4 ºC. DNA (5 nM) was incubated with the indicated concentrations of purified proteins for 15 min on ice in binding buffer (50 mM Tris-Cl pH 7.5, 125 mM NaCl, 5 mM MgCl_2_, 50 mM KCl, 5 % (v/v) glycerol, 1 mM DTT) in the presence or absence of 1 mM ATP. The DNA-protein mixture was then resolved by electrophoresis on 0.8% (w/v) agarose gels in Tris-acetate buffer for 1.5–2 h at 8 V·cm^−1^ at 4 ºC. Free and protein-bound DNA fractions were detected on a Typhoon FLA9500 scanner (Cytiva) with excitation at 532 nm using a 575-nm LP emission filter and analyzed with FIJI (*40*).

#### Single-molecule DNA loop extrusion assays

5’-Phosphorylated and 3’-biotinylated oligonucleotides complementary to naturally occurring overhangs were ligated to lambda phage DNA (λDNA; N3011, New England Biolabs) with T4 ligase. Glass slides with holes drilled for microfluidics and coverslips were treated as described (*6*). After cleaning in 1 M KOH and methanol, the glass surface was activated with 1 % (v/v) (3-aminopropyl)triethoxysilane (440140, Sigma) in 5 % (v/v) acetic acid in methanol. The glass surface was PEGylated with a 130 g·L^−1^ 1:45 mixture of biotin-PEG-SVA (MW 5,000 Da) and methyl-PEG-SVA (MW 5,000 Da; BIO-SVA_M-SVA, Laysan) in 0.1 M sodium bicarbonate buffer overnight and stored *in vacuo* at –20 ºC. Immediately prior to imaging, glass surfaces were treated with 25 mM methyl-PEG_4_-NHS (22341, ThermoFisher) in 0.1 M sodium bicarbonate. After assembly of the microfluidic device, chambers were flushed with 25 µg/mL streptavidin (189730, Merck) in rinsing buffer (50 mM Tris-HCl pH 7.5, 20 mM NaCl, 0.2 mM EDTA) and rinsed thoroughly before introducing 1–10 pM biotinylated λDNA in imaging buffer (50 mM Tris-HCl pH 7.5, 50 mM NaCl, 2.5 mM MgCl_2_, 5 % (w/v) D-glucose, 1 mM DTT, 500 nM SytoxOrange [S11368; ThermoFisher], 40 µg/mL glucose oxidase (G2133, Sigma), 15 µg/mL catalase (C1345, Sigma), 2 mM Trolox (238813, Sigma)) during continuous flow (2– 8 µL/min) using a PHD2000 syringe pump (Harvard Apparatus).

Unbound DNA molecules were flushed out with imaging buffer before introduction of one chamber volume (∼20 µL) of 0.15–2.0 nM purified condensin and 0.4–1.0 mM ATP in imaging buffer. For loop extrusion rate analyses, condensin protein concentrations were titrated down to avoid multiplex events on the same DNA molecule. In side-flow experiments, a secondary flow inlet at an angle was used with higher flow rates (15*–*25 µL/min). For TEV-cleavage experiments, 0.4 mM ATP and 0.3 µM TEV protease were added to the ATTO647N-conjugated condensin on ice immediately prior to imaging. All *Ct* condensin buffers were pre-equilibrated to ambient temperature (∼25 ºC) before protein addition, whereas *Sc* experiments were assembled on ice.

Images were acquired on a Leica GSDIM TIRF microscope system with a UPlanSApo NA 1.43 160× objective (Leica) illuminating in highly inclined and laminated optical sheet mode (HILO) with 100-ms exposure with a 532-nm laser alone, or alternating with 150-ms exposures with a 643-nm laser, all detected by an EMCCD camera (iXon Ultra 897, Andor). Overall imaging rates were 7.3 Hz for SxO-only experiments and 2.0 Hz for dual-color imaging. DNA loop extrusion by Ycs4 mutant condensin was partially acquired on a Zeiss Elyra 7 system with a Plan-Apochromat NA 1.4 63× oil objective (Zeiss) in HILO mode with 100-ms exposure with a 488-nm laser and detection with a PCO edge 4.3 CL HS sCMOS camera (PCO) at 10 Hz.

#### DNA loop extrusion image analysis

Imaging data was processed in FIJI with a custom script. SxO pixel intensity information was projected in one dimension by summing 15–21 pixels (100 nm/pixel) along the DNA double-tether trajectory for each frame and assembled into kymographs. Median background and noise were determined from 5 pixels above and below the DNA from all frames and a Gaussian-blurred (σ = 1.5) copy was generated for segmentation. The general DNA-containing area was segmented by a background + 1.5 × noise threshold and the DNA end-to-end tether length was defined as the median distance over all time points between the outer-most pixels that exceeded the median pixel intensity in the DNA-segmented area minus noise. Intensity maxima were detected per frame in the blurred copy and pixel intensity values of the 9 pixels around the maximum were summed and attributed to the “loop” region and those above or below this region as “above” or “below”. To correct for non-looped DNA within the looped region, the mean intensity value of DNA outside the “loop” region was subtracted from the loop region and assigned to the outer regions. Total intensity was normalized to 48,502 bp to convert intensity values into nucleotide lengths (movie S1). Loop extrusion rates were defined by the slope of the linear fit with the highest R^2^ of 50 consecutive data points within a user-defined time window. For *Chaetomium thermophilum* condensin loop extrusion rates, the time window for the linear fit of at least 50 consecutive data points was selected manually, due to complex loop extrusion behavior.

Because the preformed loop had often already reached its stalling size and regularly slips from its anchor point, we deduced translocation rates after direction changes from the reduction in the DNA ahead of the loop. The extension of the DNA outside of the loop was inferred from the median end-to-end tether length over the expected contour length of the total calculated base pairs outside of the loop, taking 0.41 nm·bp^−1^ to correct of 500 nM SxO intercalation (*6*). Tensions were estimated from the extension values according to the worm-like chain model (*54*), taking 42 nm as persistence length for our buffer composition.

For TEV cleavage experiments, DNA molecules were excluded when they a) did not show DNA loop formation (∼30%), b) showed loop formation but no loop rupture during imaging (∼20%), c) showed loop ruptures but the initiating condensin molecule was non-fluorescent (∼10 %), d) showed ruptures of loops initiated by a fluorescent condensin molecule but had accumulated multiple fluorescent condensins at the loop before rupture (∼15 %), 5) or when they showed ruptures of loops initiated by a fluorescent condensin molecule that had bleached before rupture (∼10 %).

Slipping and direction change events by *Ct* condensin were determined by these following criteria: A minimum change within the DNA region of 5 kbp and a minimum duration of 10 s. Reeling or slipping rates were determined from the downward or upward slope of the relevant outer region, respectively.

**Fig. S1.**
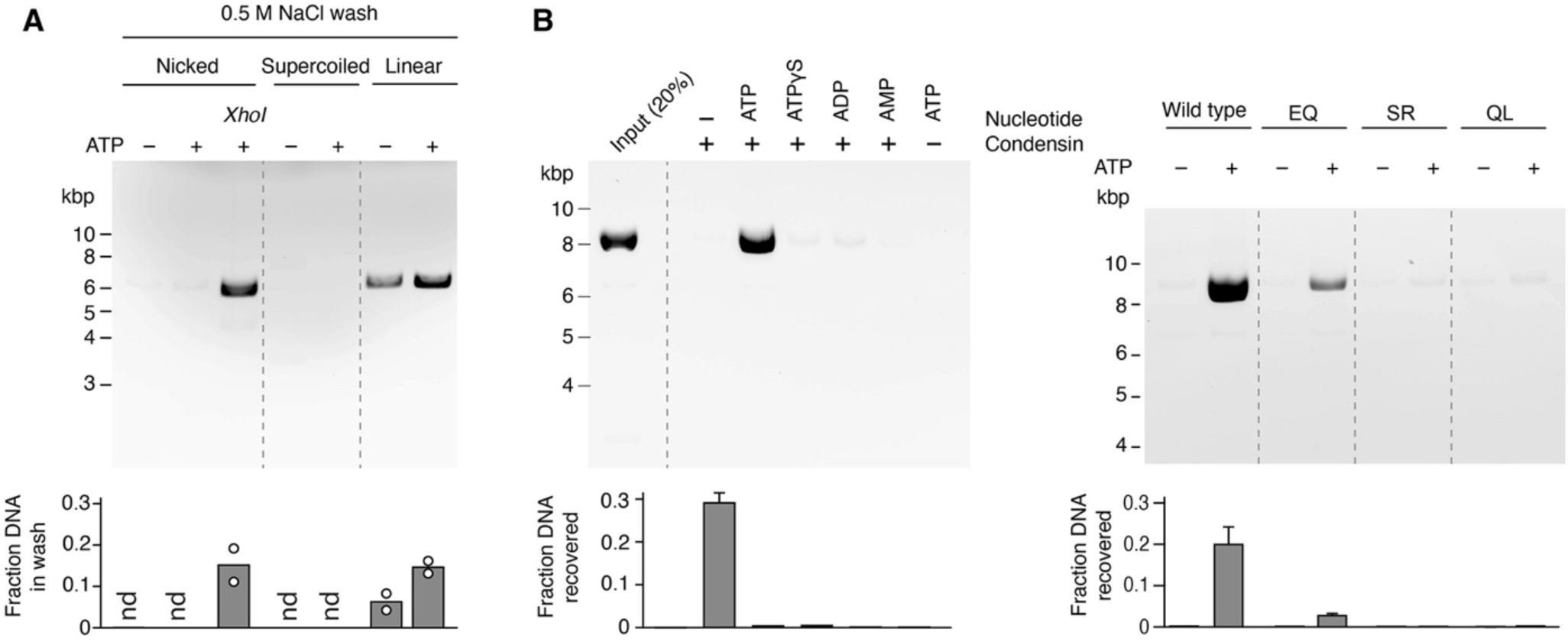
Formation of salt-resistant condensin-DNA complexes requires circular DNA and hydrolyzable ATP. (**A**) Condensin-DNA complexes were subjected to 0.5 M salt washes and the released DNA was resolved by agarose gel electrophoresis and quantified after ethidium bromide staining (mean, *n* = 2; nd = not determined). Remaining salt-resistant DNA was then eluted and is shown in Fig. 1B. (**B**) Formation of salt-resistant condensin-DNA complexes by wild-type condensin in the presence of different nucleotides (left) and by wild-type, EQ Walker B mutant (Smc2_E1113Q_; Smc4_E1352Q_), SQ signature motif mutant (Smc2_S1085R_; Smc4_S1324R_) or QL Q-loop mutant (Smc2_Q147L_; Smc4_Q302L_) in the presence of ATP (right) as quantified in Fig. 1B (mean ± s.d., *n* = 3).

**Fig. S2.**
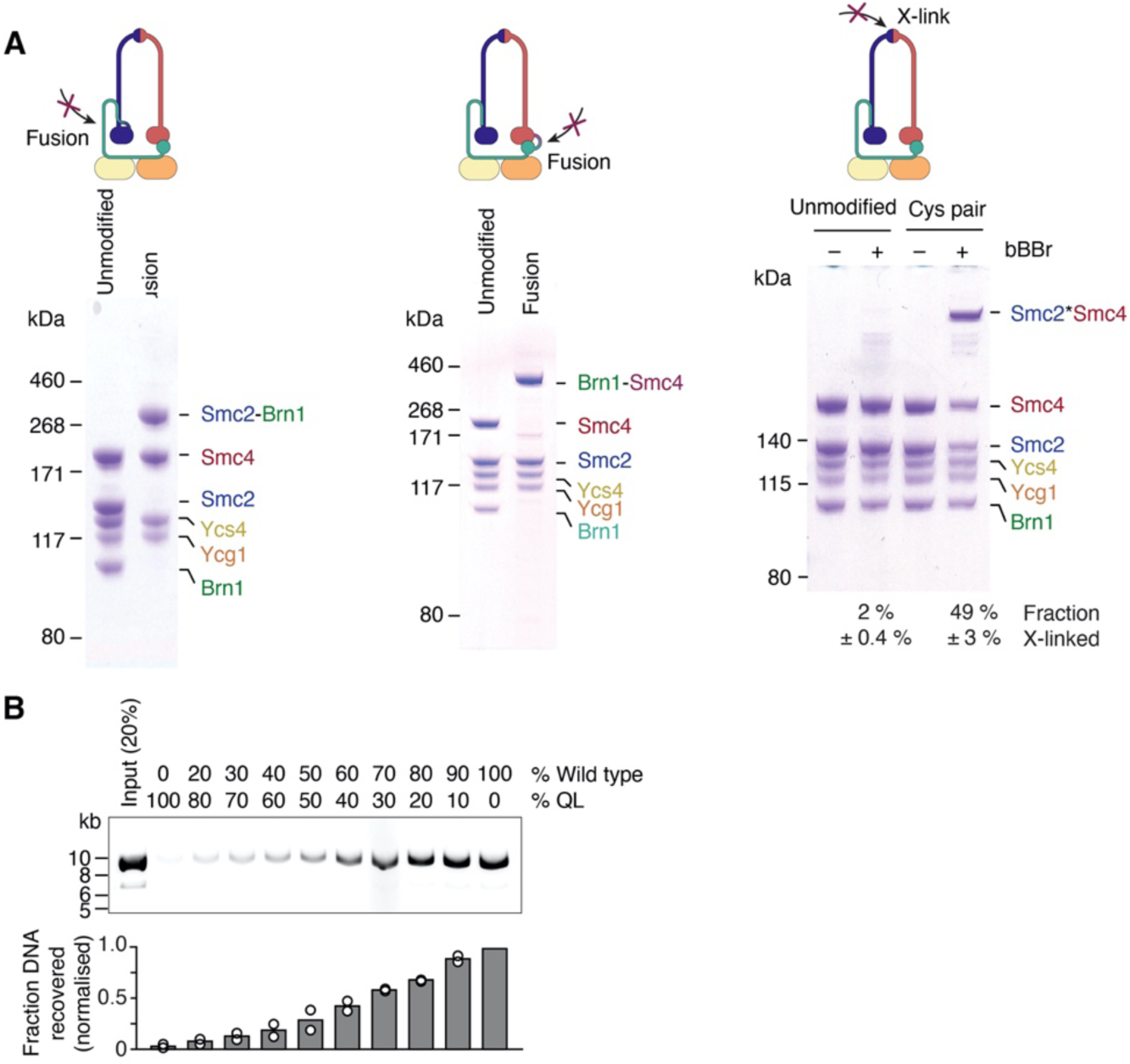
Closure of SMC-kleisin ring interfaces. **(A)** SDS-PAGE of purified unmodified, Smc2−Brn1 fusion, Brn1−Smc4 fusion, or cross-linkable hinge (Smc2_K639C_; Smc4_V721C_) condensin complexes after staining with Coomassie Brilliant Blue. The fraction of cross-linked species was estimated by densitometry (mean ± s.d., *n* = 3). **(B)** Formation of salt-resistant condensin-DNA complexes using a mixture of wild-type and QL catalytic mutant (Smc2_Q147L_; Smc4_Q302L_) condensin as in Fig. 1B. The graph shows the quantitation of salt-resistant DNA fractions for each ratio after ethidium bromide staining (mean, *n* = 2).

**Fig. S3.**
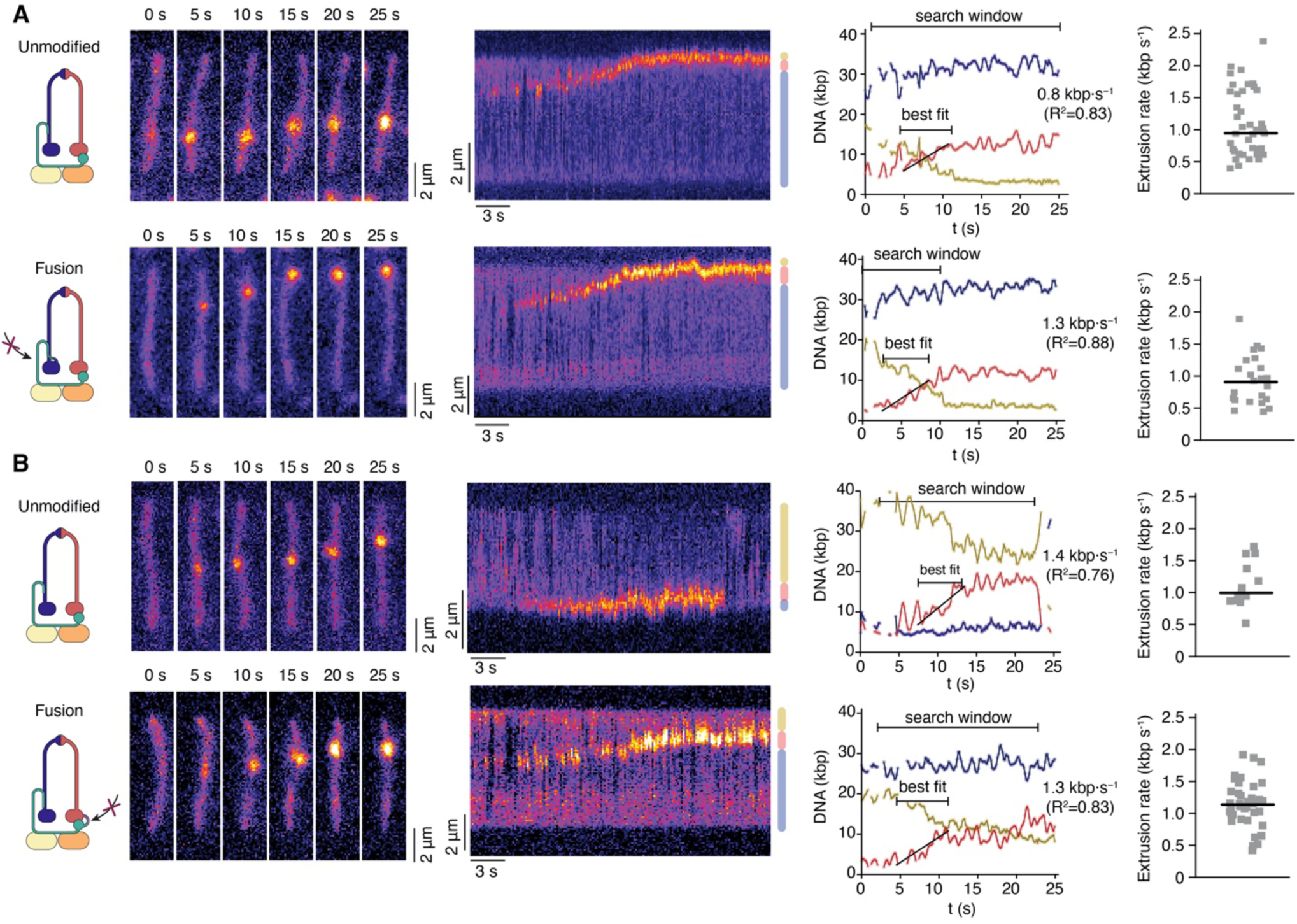
DNA loop extrusion by condensin with fused SMC-kleisin interfaces. (**A**) Stills and kymographs of surface-tethered λ-phage DNA (48.5 kbp) stained with SxO and incubated with unmodified or Smc2−Brn1 fused condensin. Graphs show quantitation of DNA in (red), above (yellow) and below (blue) the loop. A user-defined search window was selected in which the slope of the best linear fit of 50 consecutive data points was taken as loop extrusion rate. Individual loop extrusion measurements and the median (line) are plotted. (**B**) As in (A) for Brn1−Smc4 fusion condensin. See also movie S1.

**Fig. S4.**
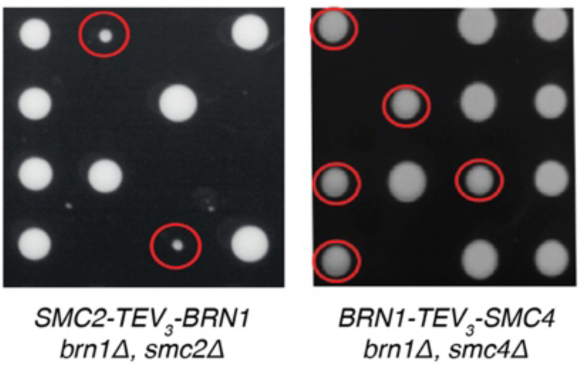
*In vivo* functional analysis of SMC-kleisin fusion proteins. Tetrad dissection of diploid yeast strains after 3 days at 25 °C. Colonies of cells expressing Smc2–Brn1 or Brn1–Smc4 fusion proteins as their sole source of Brn1 and Smc2 or Smc4, respectively, are indicated by circles

**Fig. S5.**
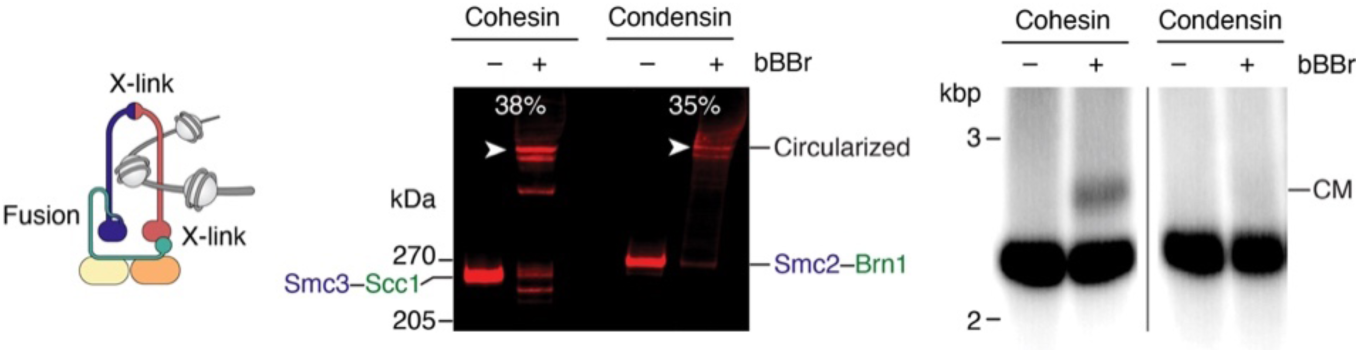
*Ex vivo* circularization of cohesin or condensin SMC-kleisin rings on mini-chromosomes. Lysates from budding yeast strains containing a circular 2.3-kbp *TRP1-ARS1-CEN4* mini-chromosome and either covalently circularizable cohesin (Smc1_G22C, K639C_; Smc3_E570C_–Scc1_A547C_-HA_6_ fusion) or condensin (Smc4_V721C, R1417C_; Smc3_K639C_–Brn1_K709C_-HA_6_ fusion) were subjected to anti-HA epitope immunoprecipitation and treated with bBBr cross-linker. Protein-DNA complexes were eluted in 1% SDS at 65 °C and analyzed by anti-HA fluorescent Western blotting to resolve protein cross-links (left) or by Southern blotting to detect mini-chromosomes and their covalent monomeric catenanes (CM) with circularized SMC-kleisin rings (right). The fractions of circularized cohesin or condensin are indicated.

**Fig. S6.**
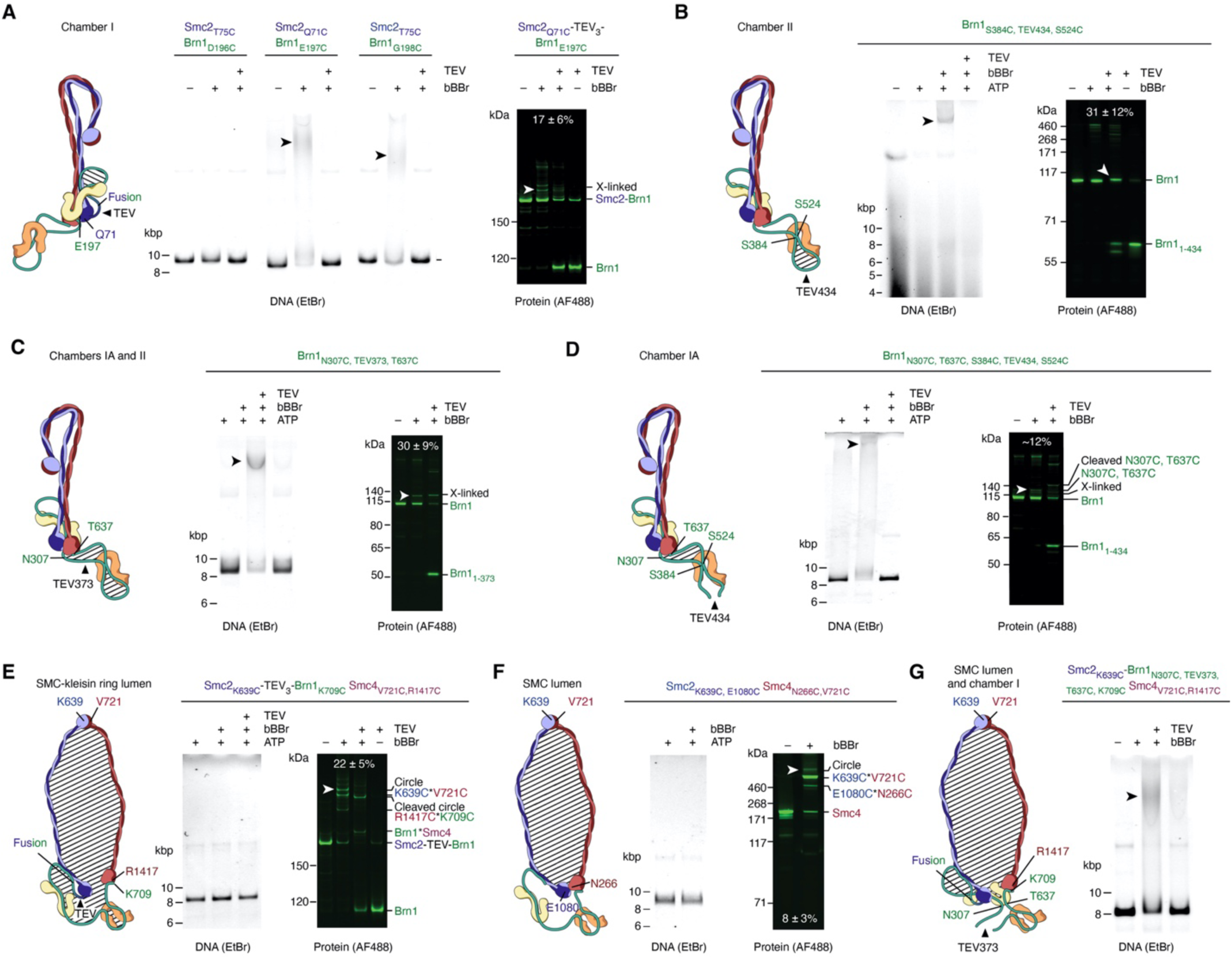
Effect of cross-linking different condensin compartments on retaining protein-DNA catenanes after protein denaturation. (**A**) Circularization of chamber I in condensin-DNA complexes shown in Fig. 2B. In-gel fluorescence detection of DNA after SDS protein denaturation, agarose gel electrophoresis and ethidium bromide staining (left) and of Smc2–TEV_3_–_AF488_Brn1 protein after SDS-PAGE to determine cross-linking efficiency (mean ± s.d., *n* = 4) and confirm efficient TEV cleavage. Catenanes and cross-linked species are highlighted with arrowheads. (**B**) Circularization of chamber II shown in Fig. 2C by cross-linking Brn1_N307C, T637C_ as in (A). Cross-linking efficiency is indicated (mean ± s.d., *n* = 9). **(C)** Circularization of combined chambers IA and II as in (A). Cross-linking efficiency is indicated (mean ± s.d., *n* = 5). (**D**) Selective circularization of chamber IA by combined cross-linking Brn1_N307C, T637C_ and Brn1_S384C, S524C_ with simultaneous opening of chamber II by TEV434 cleavage as in (A). (*n* = 2) (**E**) Circularization of the SMC-kleisin ring using cysteine cross-linking of hinge (Smc2_K639C_; Smc4_V721C_) and Smc4–Brn1 (Smc4_R1417C_; Brn1_K709C_) interfaces in combination with Smc2−TEV–Brn1 fusion as in (A). Cross-linking efficiency is indicated (mean ± s.d., *n* = 5). (**F**) Circularization of the SMC lumen using cysteine cross-linking of hinge (Smc2_K639C_; Smc4_V721C_) and engaged ATPase domains (Smc2_E1080C_; Smc4_N266C_) in the presence of ATP. Cross-linking efficiency is indicated (mean ± s.d., *n* = 4). (**G**) Circularization of combined SMC lumen and chamber I using cysteine cross-linking of hinge (Smc2_K639C_; Smc4_V721C_), Smc4–Brn1 (Smc4_R1417C_; Brn1_K709C_) and Brn1_N307C, T637C_ and opening combined chambers IA and II using TEV373 cleavage as in (A).

**Fig. S7.**
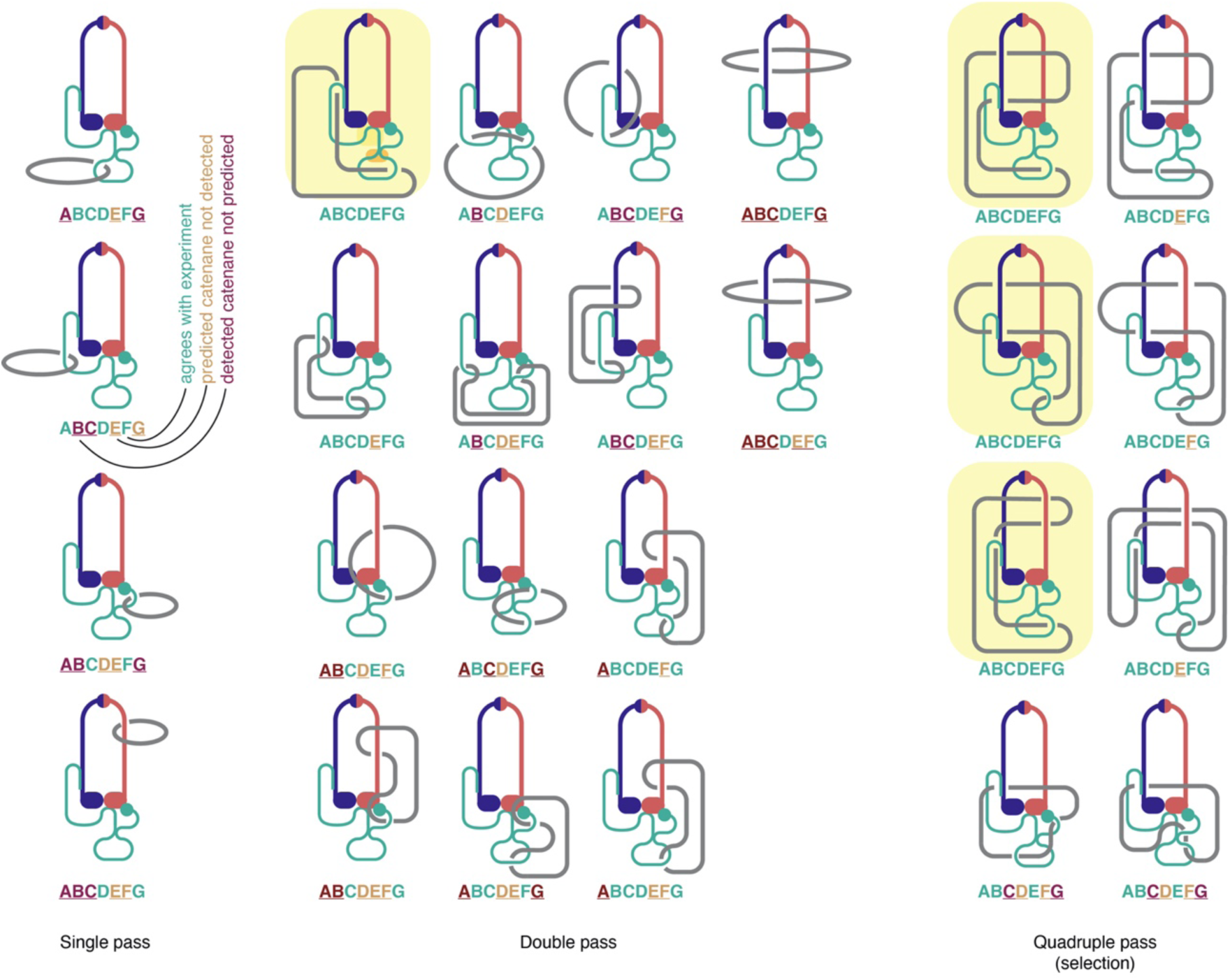
Overview of possible condensin-DNA topological configurations. Schematic depiction of theoretical configurations between a condensin complex with four compartments and circular DNA. Up to two DNA segments are allowed in the SMC lumen. Four unique configurations exist for a single pass of the DNA (left) and seven combinations for a double pass, with each two ways to connect the two DNA segments (center). A selection of unique configurations for a quadruple pass is also shown (right). Letters (A–G) below each configuration refer to the topological experiments in fig. S6A–G, with colors marking agreement (green), or disagreement (orange and red) with the experimental results. Configurations that are in line with all topological experiments are highlighted.

**Fig. S8.**
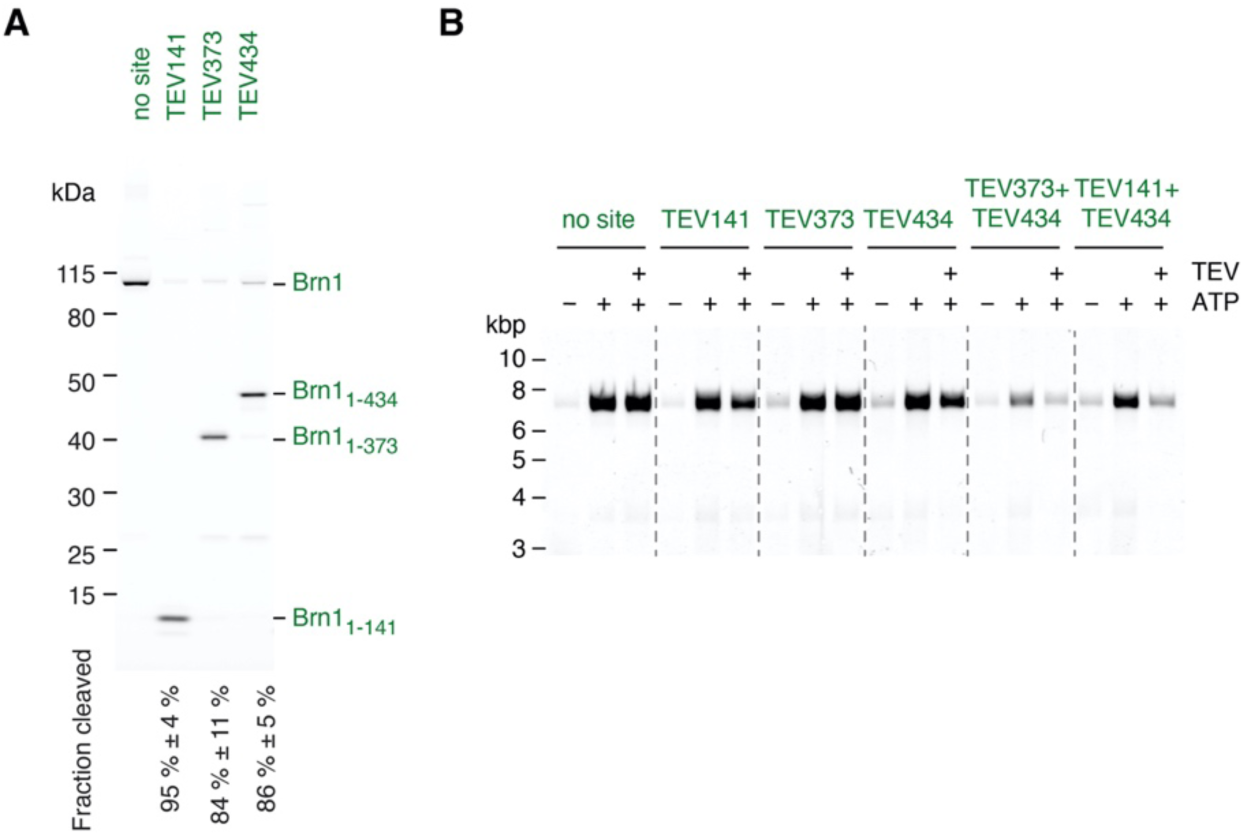
Effect of TEV cleavage of kleisin chambers on salt-resistant condensin-DNA complexes. (**A**) SDS-PAGE of condensin complexes with indicated TEV cleavage sites and quantitation of cleavage efficiencies by in-gel fluorescence detection of _AF488_Brn1 after TEV incubation (mean ± s.d., *n* ≥ 6). (**B**) Fluorescence detection of salt-resistant condensin-DNA complexes with TEV-cleavable Brn1versions after agarose gel electrophoresis and ethidium bromide staining.

**Fig. S9.**
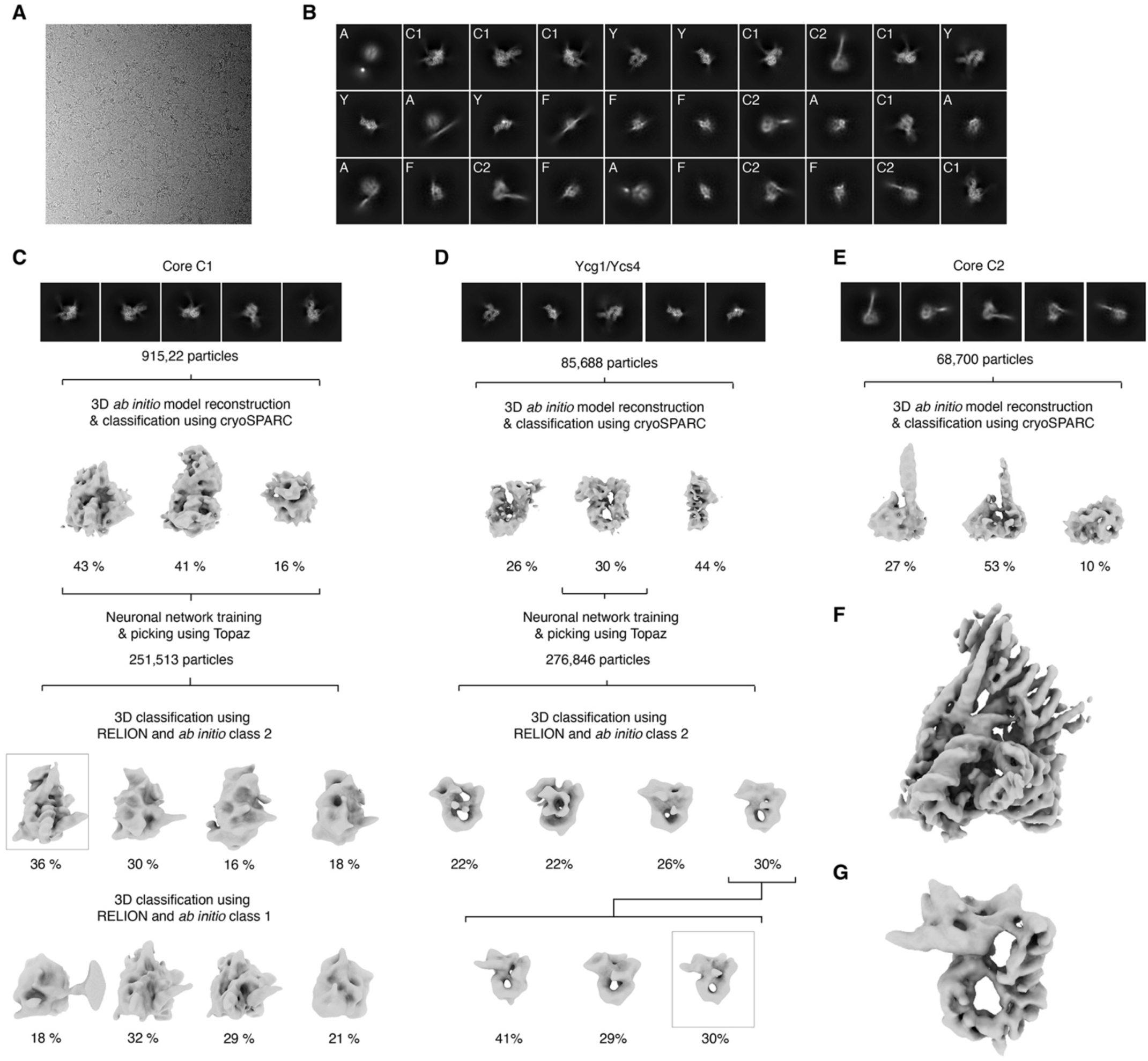
Initial cryo-EM model generation. (**A**) Representative micrograph of condensin-DNA particles. (**B**) Representative 2D classes of the data set showing particles in conformations C1 (core 1), C2 (core 2), Y (Ycg1/Ycs4) or F (fragments). (**C–E**) Workflow of initial data processing of C1, Y or C2 particles, respectively. (**F, G**) 3D-refined maps of core or peripheral subcomplexes, respectively, bound to DNA. The maps were used for further processing as depicted in fig. S10.

**Fig. S10.**
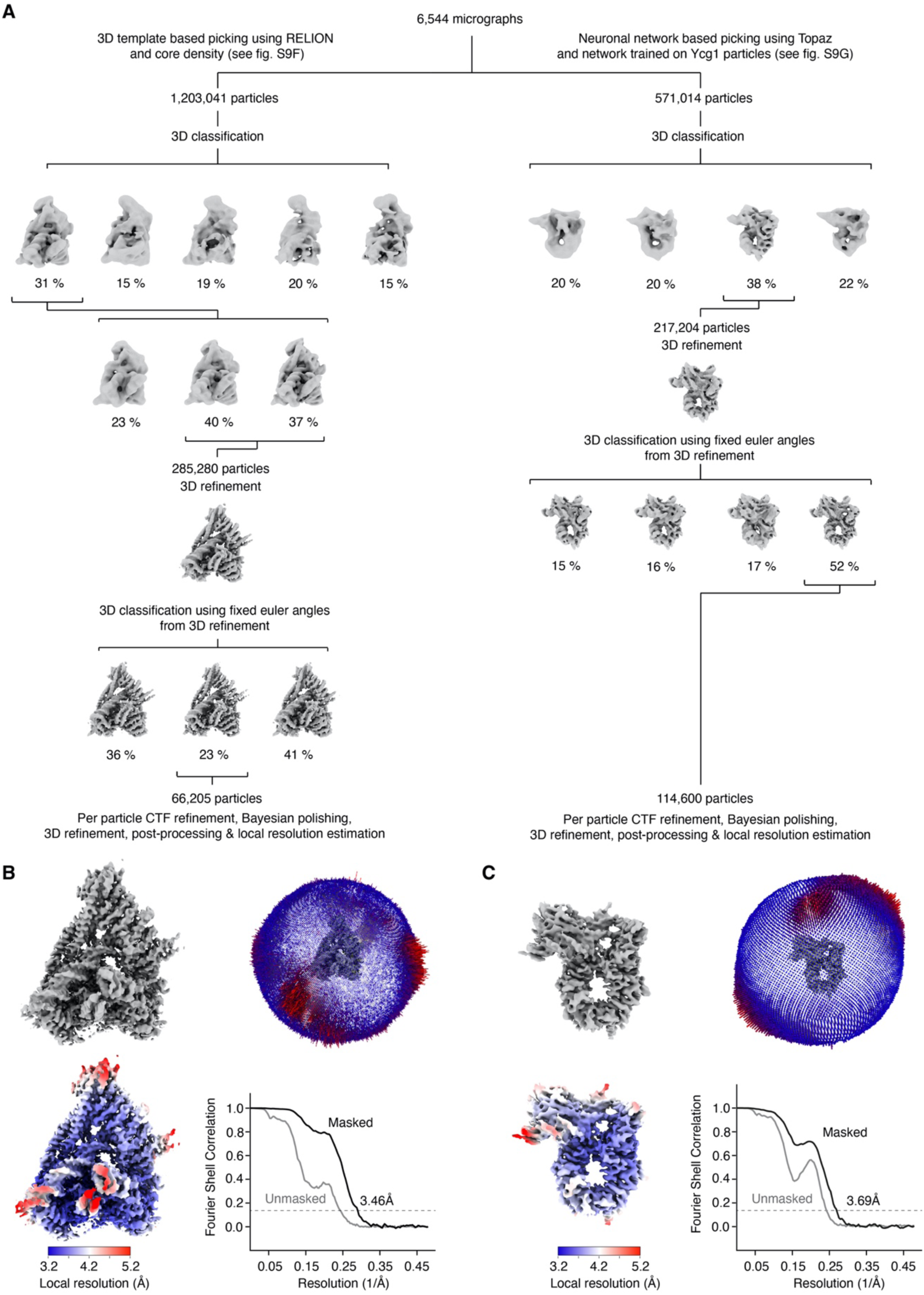
Cryo-EM refinement procedures. (**A**) Processing of the core (left) and peripheral (right) subcomplexes bound to DNA. Particles were initially processed and preliminary angles and translations were assigned as described in fig. S9C and D. 3D classifications, 3D refinements, per particle CTF refinements, Bayesian polishing, post-processing and local resolution estimations resulted in the final volumes, angular distribution plots, local resolution maps and FSC curves shown in (**B**) and (**C**).

**Fig. S11.**
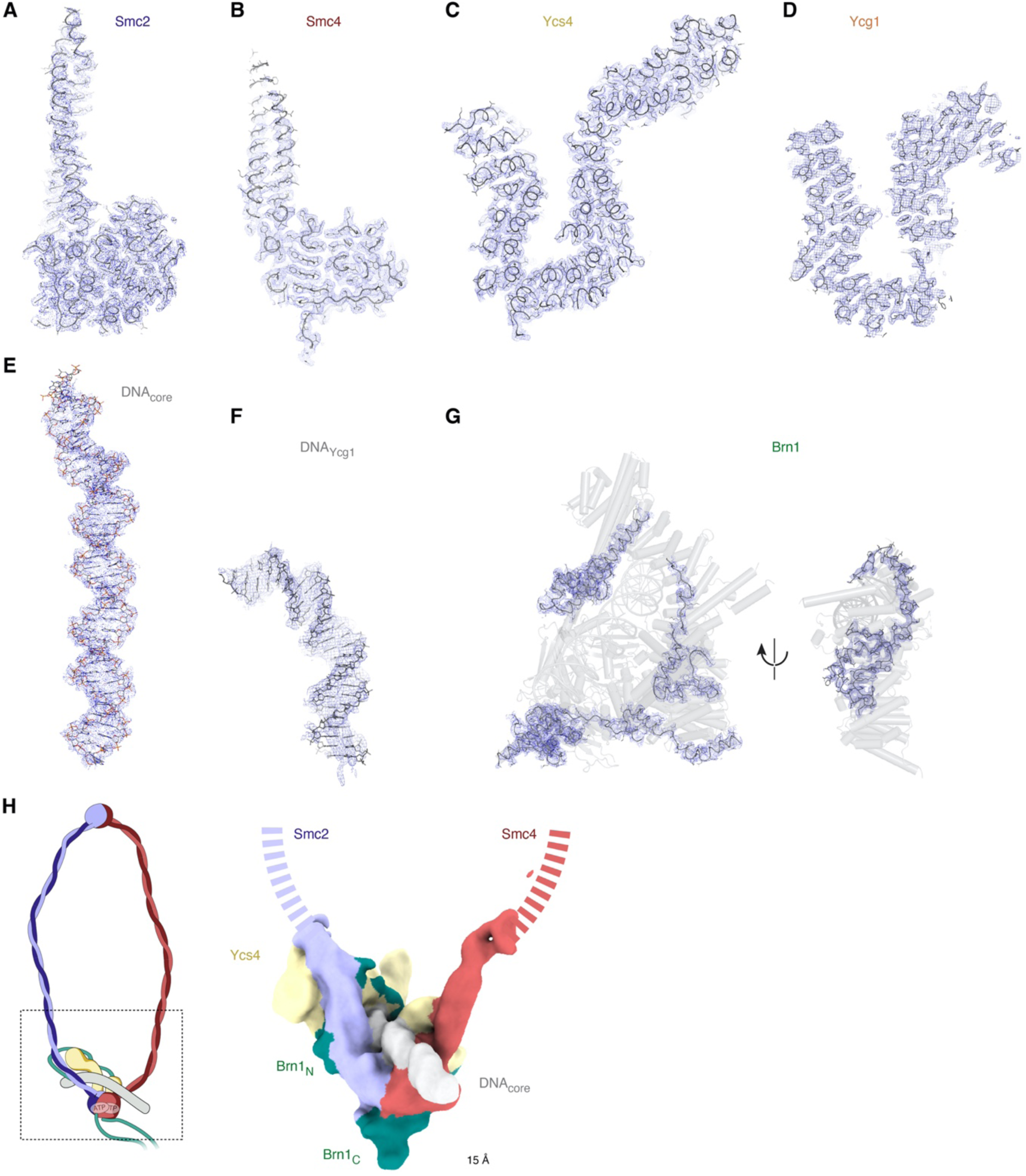
Electron density maps of individual subunits. Electron density maps showing (**A**) Smc2, (**B**) Smc4, (**C**) DNA, (**D**) Ycs4, (**E**) DNA bound to the core module, (**F**) DNA bound to the periphery module, DNA bound to peripheral module, or (**G**) the ordered Brn1 segments. (**H**) The trajectory of the open Smc2 (blue) and Smc4 (red) coiled coils is clearly discernable in a 15-Å resolution map of the core module.

**Fig. S12.**
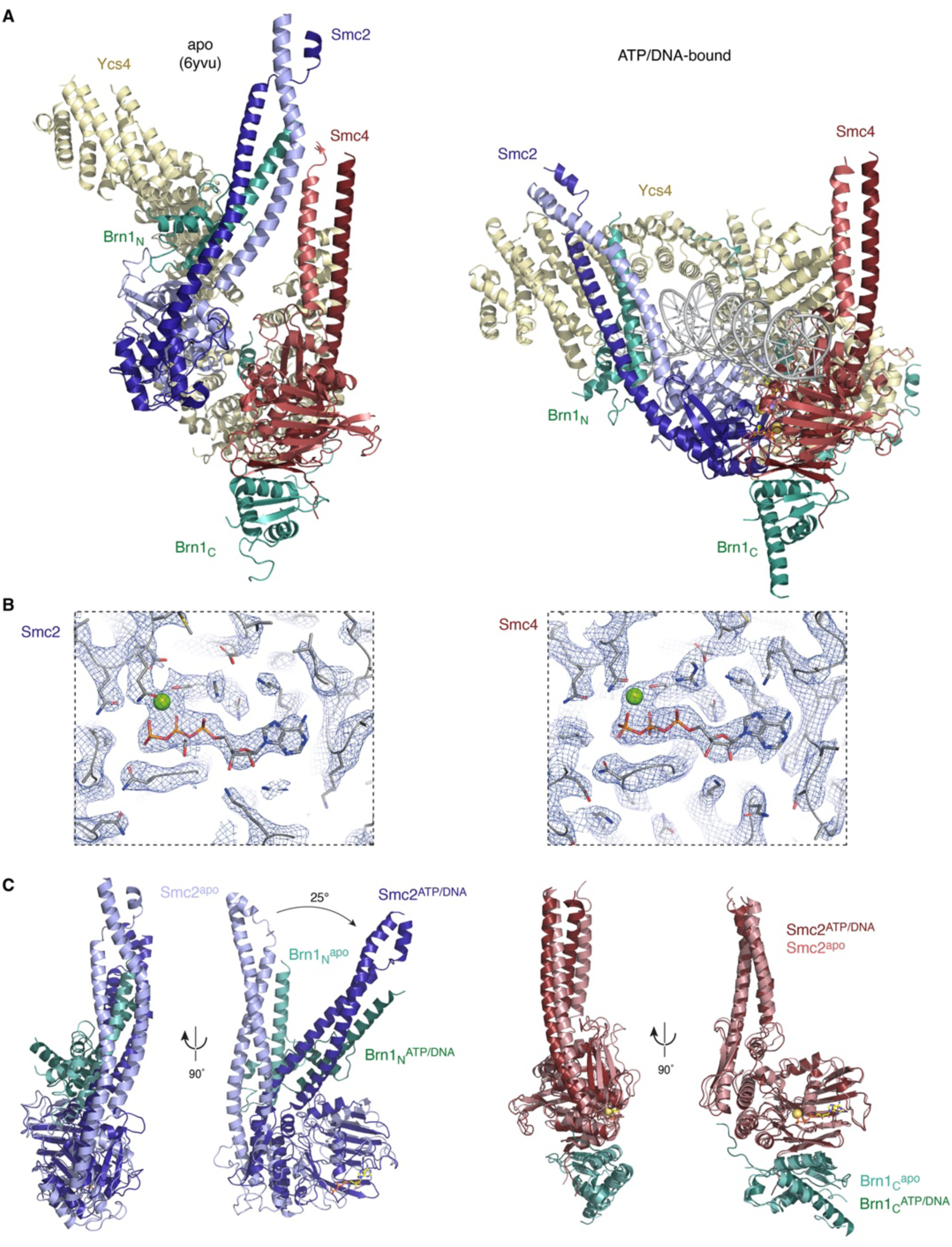
Conformational changes of Smc2 and Smc4 heads upon ATP binding. Comparison of ATP/DNA-bound (dark) to apo condensin (pdb: 6yvu, light) structures. Overlays between Smc2_head_ and Smc4_head_ structures aligned to their respective RecA lobes highlight the coiled-coil movement induced by ATP binding.

**Fig. S13.**
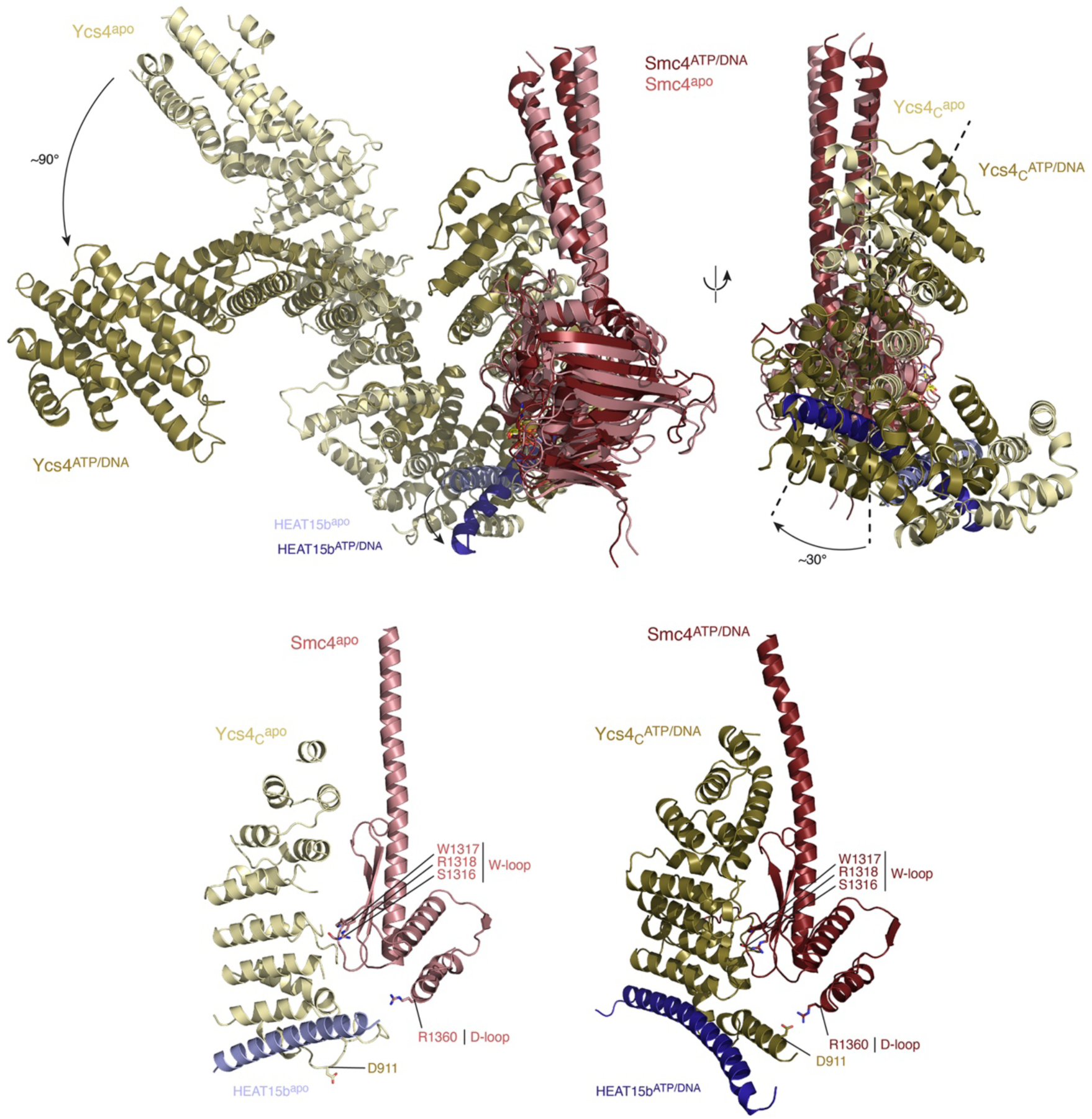
Ycs4 conformational changes upon ATP binding. (**A**) Overlay of ATP/DNA-bound (dark) and apo condensin (pdb: 6yvu, light) structures aligned to the Smc4 (red) RecA lobe. Swivel motions of the Ycs4_N_ (left) and Ycs4_C_ (right) are indicated by arrows. Zoom-in views (bottom) highlight the interface between Ycs4_C_ and the Smc4 W-loop region and the relocation of Ycs4 HEAT repeat 15b helix (blue). (**B**) Zoom-in view of the Ycs4_N_ interface with the three-helix bundle formed by the Smc2_neck_ coiled coil and Brn1_N_.

**Fig. S14.**
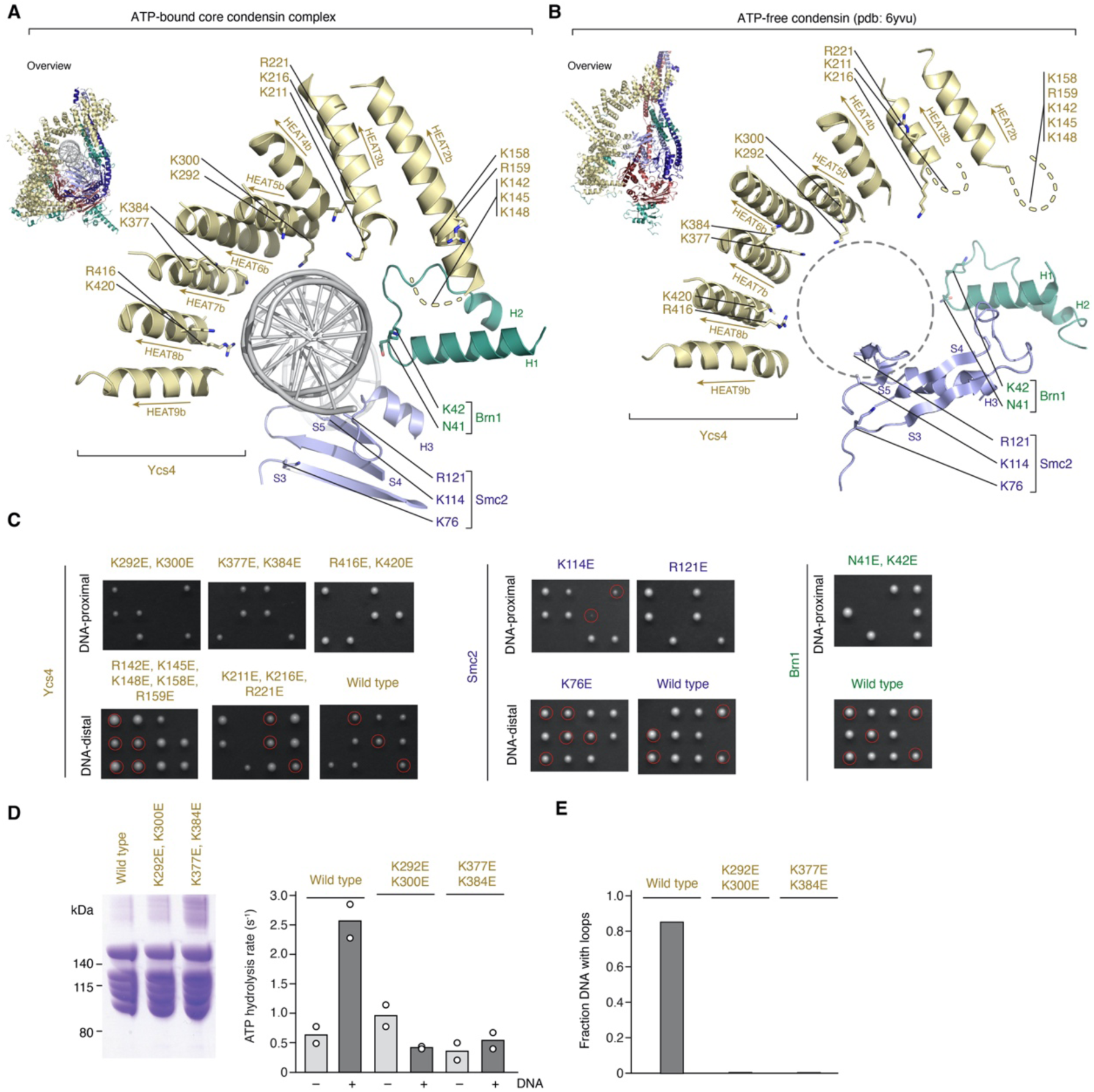
Kleisin chamber I DNA backbone contacts. (**A**) Cartoon representation of the DNA-binding chamber I in the ATP-bound state. Sidechains of basic residues of Ycs4 (yellow), Smc2 (blue) and Brn1 (green) that contact the DNA backbone (DNA-proximal) or face away from the DNA (DNA-distal) and were selected for charge-reversal mutations are highlighted. (**B**) As in (A) for the ATP-free apo state. (**C**) The ability of charge-reversal mutations in the indicated condensin subunits to support proliferation of yeast tested by tetrad dissection. Progeny of spores with a modified allele as the sole copy are highlighted by circles. (**D**) SDS-PAGE of purified *Sc* condensin holo-complexes with Ycs4 mutations and ATP hydrolysis assay in the absence and presence of 6.4-kbp nicked circular DNA (25 nM; mean, *n* = 2). (**E**) Fraction of DNA molecules that display loops in single-molecule experiments with wild-type or Ycs4 mutant condensin complexes; *n* = 88, 140 and 62 DNA molecules scored.

**Fig. S15.**
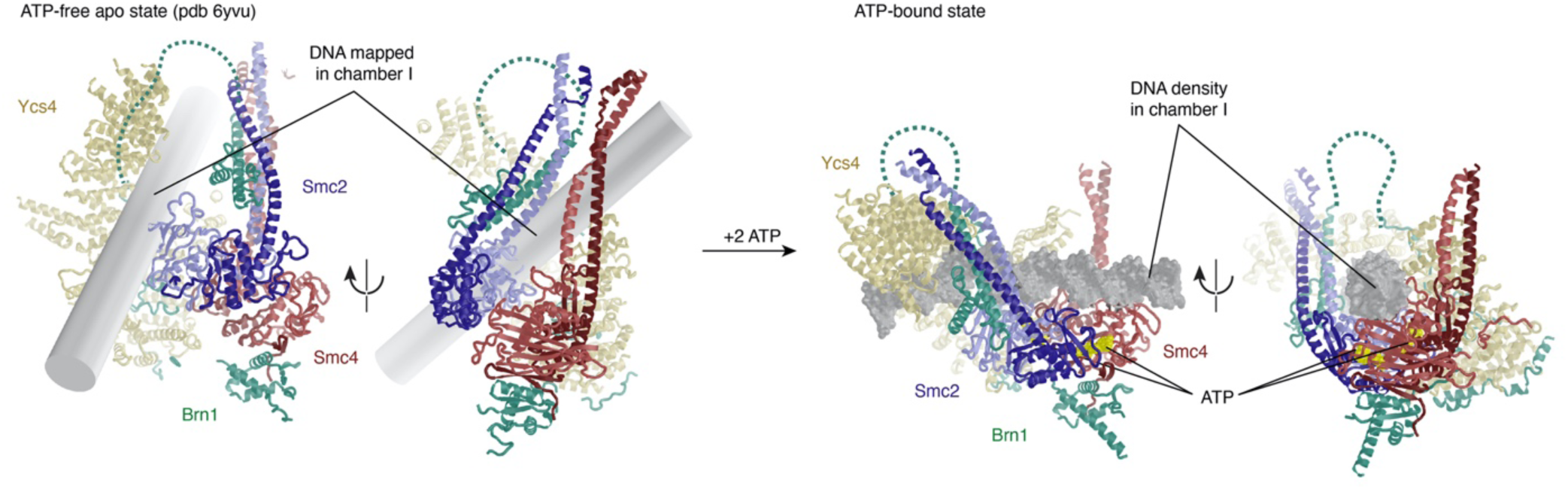
ATP binding feeds DNA between the disengaged Smc2–Smc4 coiled coils. Cartoon representation of DNA mapped into chamber I of the ATP-free apo condensin and of DNA observed in chamber I of the ATP-bound complex.

**Fig. S16.**
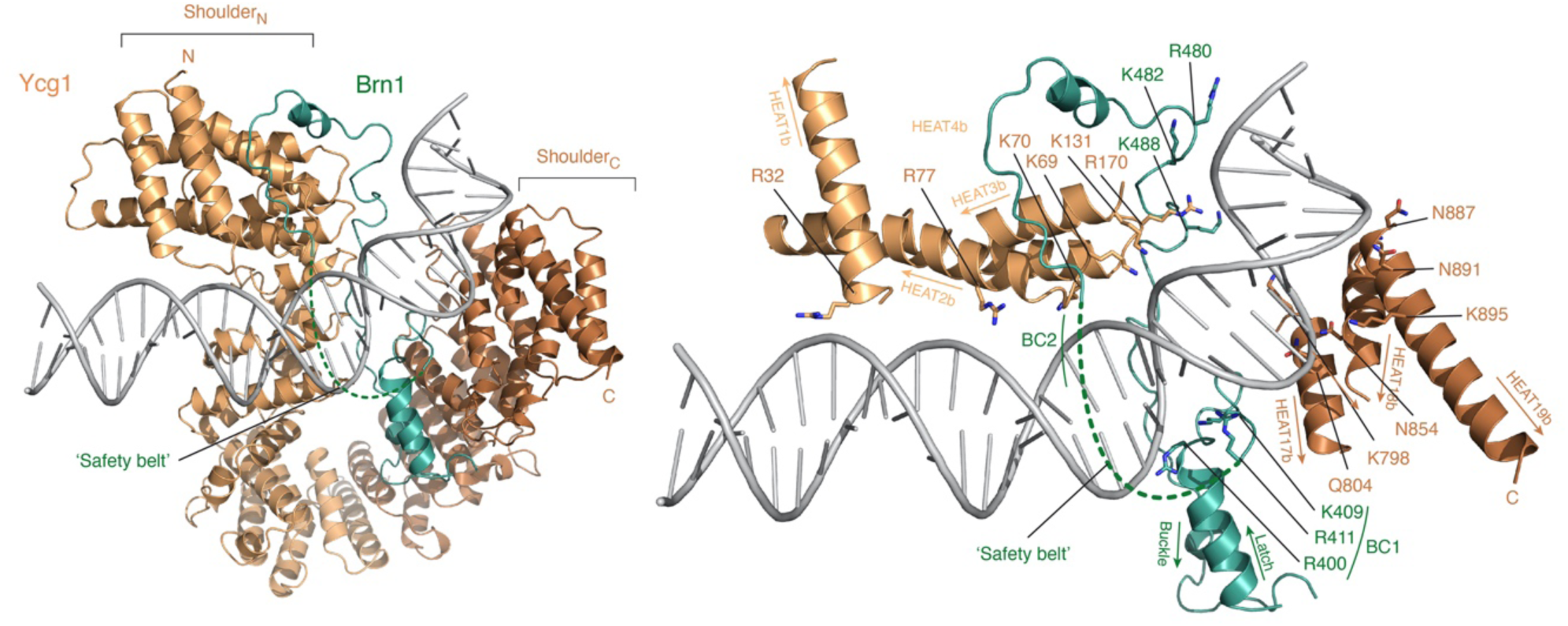
Kleisin chamber II DNA backbone contacts. Cartoon representation of the DNA-bound chamber I. Sidechains of basic residues of Ycg1 (orange) and Brn1 (green) that contact the DNA backbone are highlighted.

**Fig. S17.**
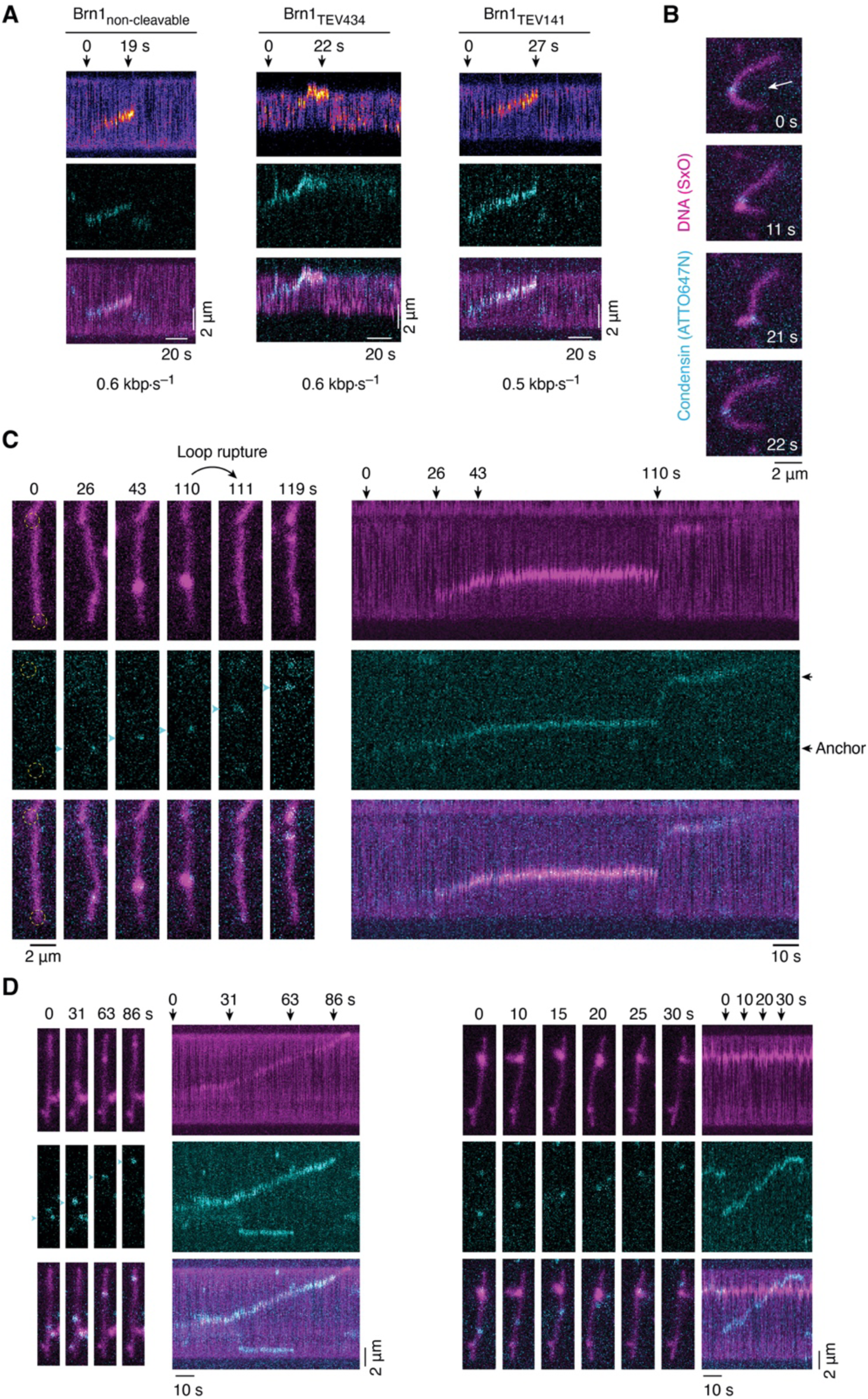
Opening of kleisin chambers I or II by TEV cleavage during DNA loop extrusion. (**A**) Kymographs of DNA loop extrusion on double-tethered λ-DNA molecules by condensin variants in presence of TEV protease and 0.4 mM ATP as depicted by stills in Fig. 4A and movies S3–S5. Extrusion rates determined as in fig. S3 using a linear fit to 20 consecutive timepoints are indicated. (**B**) Rupture of a DNA loop formed by non-cleavable ATTO647N-condensin in side-flow experiments. Arrow indicates direction of buffer flow. (**C**) Stills and kymograph of DNA loop extrusion by condensin Brn1_TEV434_ in presence of TEV protease showing continued unidirectional translocation with minimal loop extrusion after loop rupture. (**D**) Stills and kymograph of condensin Brn1_TEV434_ treated with TEV protease translocating on DNA without expanding the loop size.

**Fig. S18.**
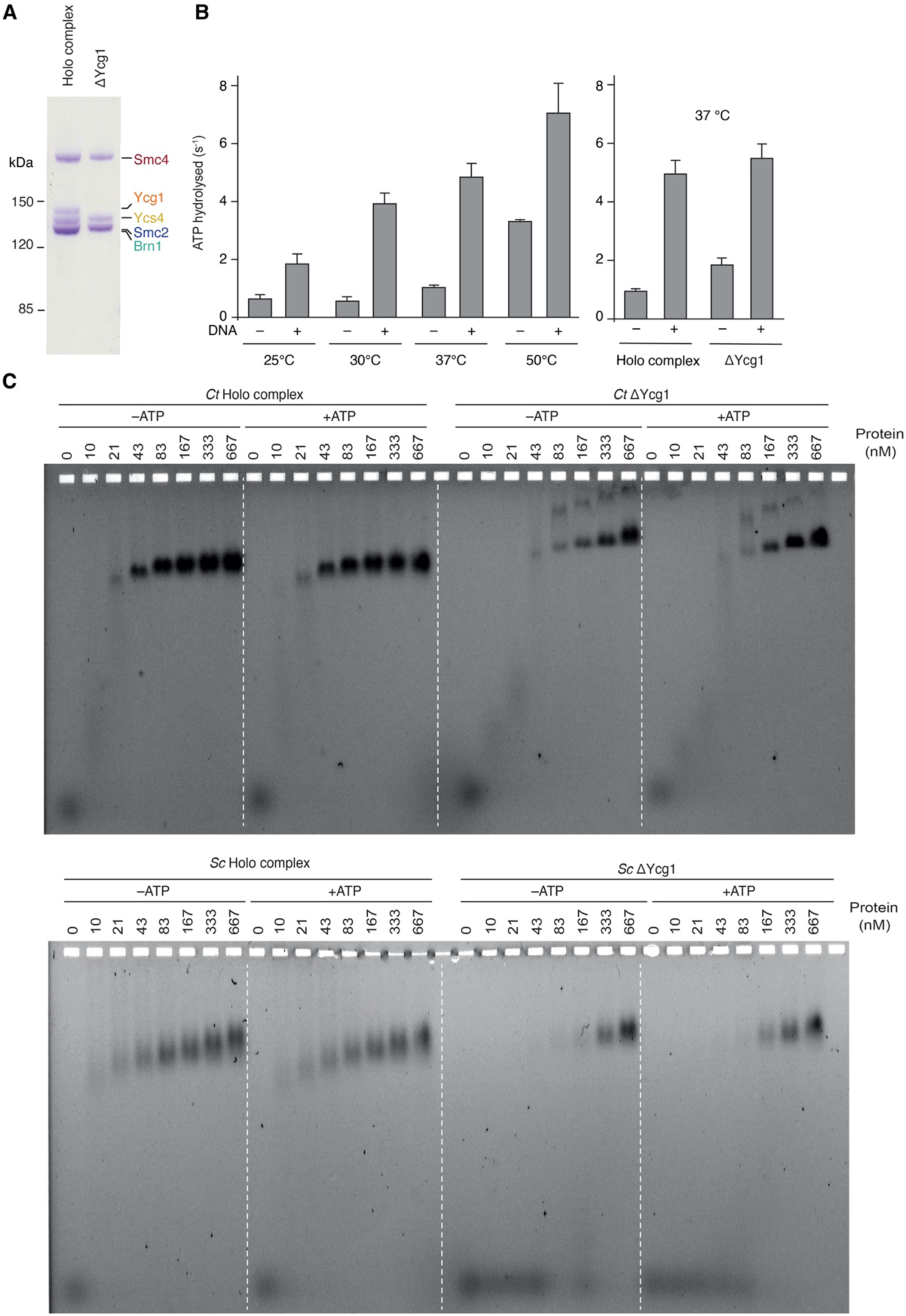
Characterization of the *Ct* condensin complex. (**A**) SDS PAGE analysis of purified *Ct* holo and ΔYcg1 condensin complexes after staining with Coomassie Brilliant blue. (**B**) ATP hydrolysis rates (molecules ATP per molecule condensin and second) at different temperatures measured for *Ct* holo condensin (0.5 µM) in the presence or absence of DNA (25 nM 6.4 kbp) and comparison of ATP hydrolysis rates for *Ct* holo and ΔYcg1 condensin (at 37 °C). (**C**) Electrophoretic mobility shift assay of a 6-carboxyfluorescin labeled 50-bp dsDNA (5 nM) at increasing concentrations of *Ct* or *Sc* holo and ΔYcg1 condensin in presence or absence of ATP (1 mM).

**Fig. S19.**
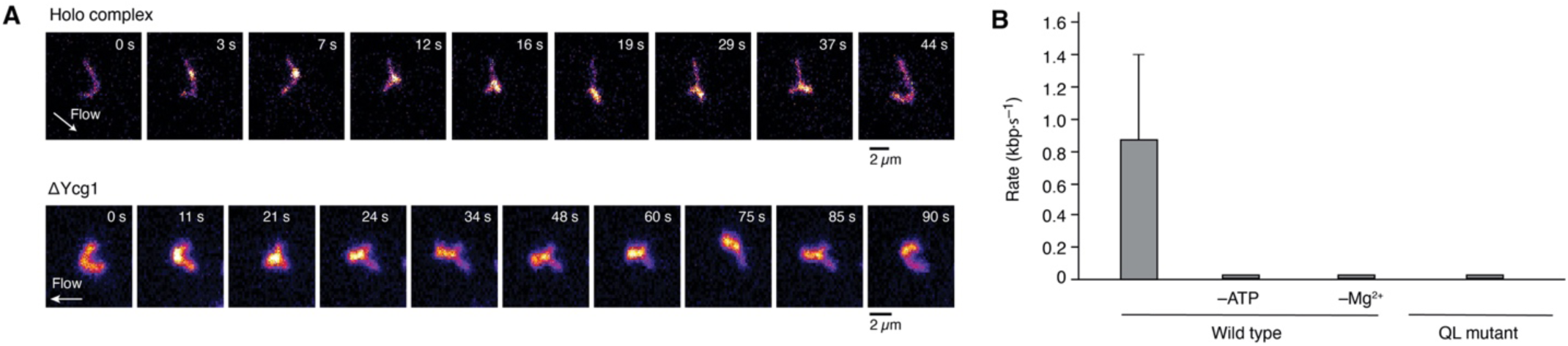
DNA loop formation by *Ct* condensin complexes. (**A**) Example image sequence of loop extrusion events by Ct holo and Ycg1 complex (below) where the extruded loop is stretched out by buffer flow approximately perpendicular to the DNA axis. (**B**) Average DNA loop extrusion rates of wild-type *Ct* condensin under various condition and an ATP-binding deficient mutant (QL; Smc2_Q147L_, Smc4_Q421L_) (mean ± s.d.; *n* = 55, 62, 56 and 60 DNA molecules measured).

**Fig. S20.**
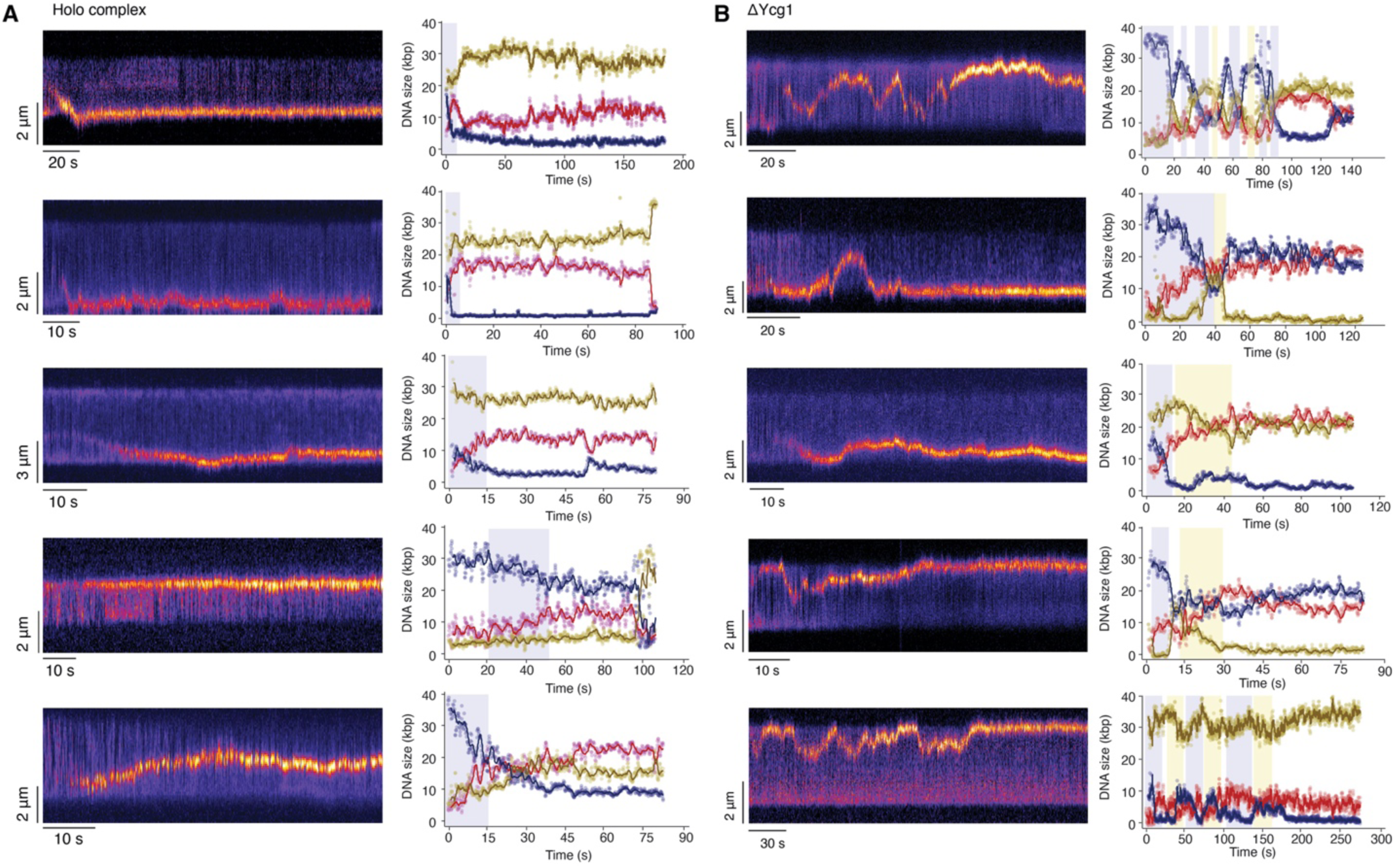
Additional examples of DNA loop extrusion events by *Ct* holo and ΔYcg1 condensin. Kymographs of DNA loop extrusion by *Ct* holo (**A**) and ΔYcg1 (**B**) condensin and fluorescence intensity plots of the DNA regions above (yellow) and below (blue) the extruded DNA loop (red). Shaded regions indicate an increase in the DNA loop and concurrent decrease of the DNA region above (yellow) or below (blue) the loop.

**Fig. S21.**
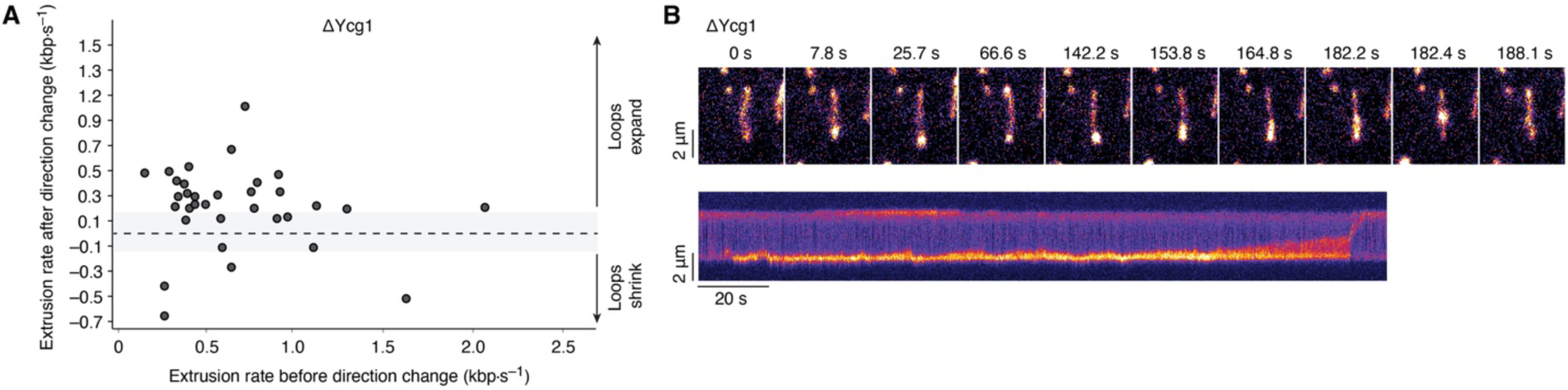
Direction change and Z-loop formation by *Ct* ΔYcg1 condensin. (**A**) Scatter plot of DNA loop extrusion rates before (x-axis) and after (y-axis) *Ct* ΔYcg1condensin switches direction. (**B**) Example image sequences and kymographs of Z-loop formation by *Ct* ΔYcg1 condensin.

**Fig. S22.**
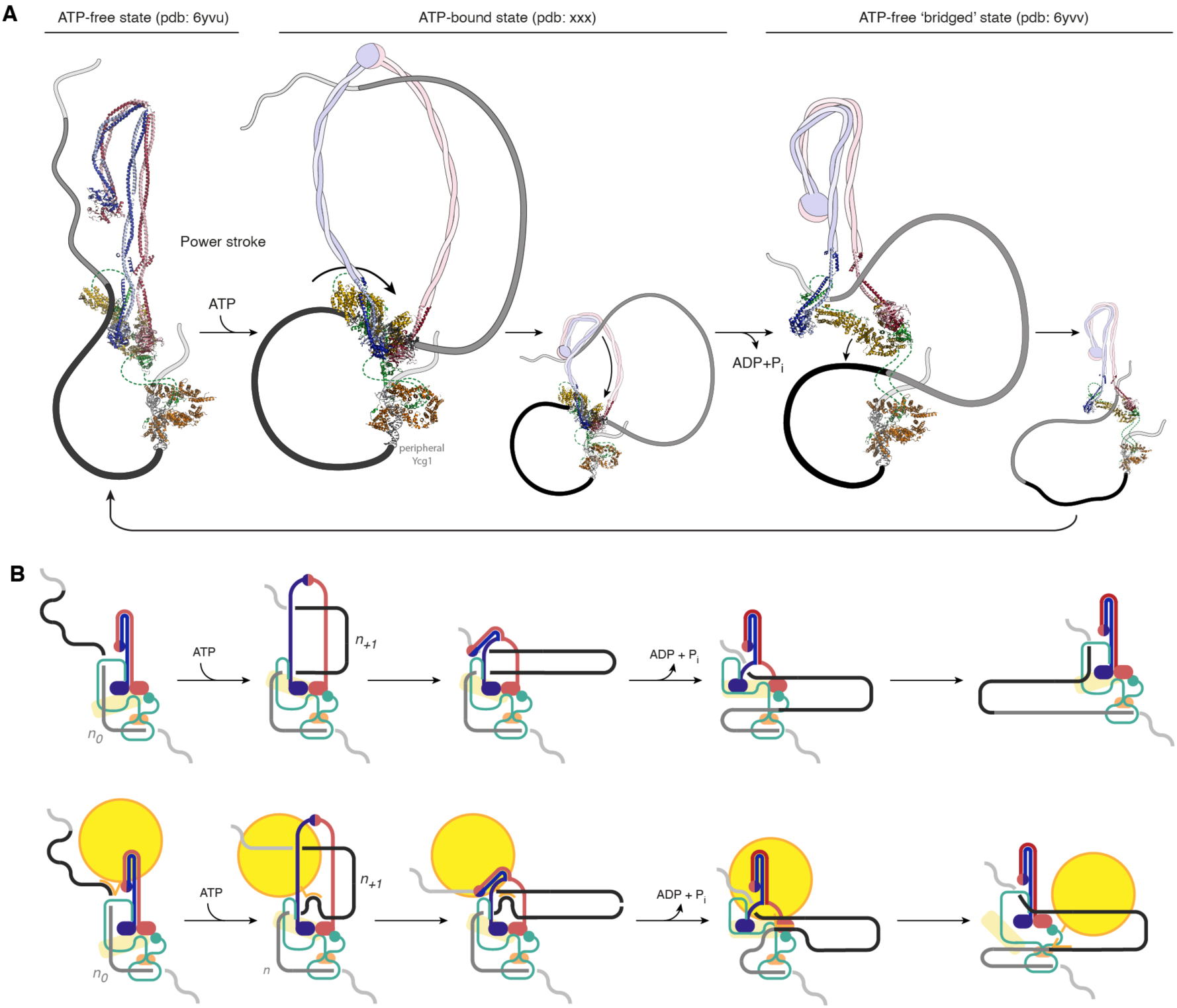
Models of the hold-and-feed mechanism. (**A**) Structural and (**B**) schematic representation of condensin and DNA topology. A pre-existing DNA loop (*n*_*0*_) is extended by a DNA loop (*n*_*+1*_) that is fed from chamber I into the coiled-coil lumen. For details see **Fig. 6**. Structural evidence for the conformation of different states of the *S. cerevisiae* condensin holo complex are shown as cartoons. Bypass of large tethered obstacles without the large obstacle entering the coiled-coil lumen are shown for the schematic representation. Opening of the Ycs4–Smc4 interface enables passage of the tether without disengagement of the SMC-kleisin ring interfaces.

**Table S1.** Cryo-EM statistics.

| **Complex** | **Sc Condensin/DNA** | **Sc Condensin/DNA** |
| --- | --- | --- |
| Subcomplex | Core/DNA | Ycg1/DNA |
| EMDB | EMD-xxx | EMD-xxx |
| PDB | xxx | xxx |
| Data collection and processing | Titan Krios | Titan Krios |
| Magnification | ×130,000 | ×130,000 |
| Voltage (kV) | 300 | 300 |
| Camera | GIF-K2 counting | GIF-K2 counting |
| Total electron exposure (e-/Å<sup>2</sup>) | 40.03 | 40.03 |
| Exposure rate (e-/pix/s) | 4.8 | 4.8 |
| Number of frames per movie | 40 | 40 |
| Automation software | SerialEM | SerialEM |
| Defocus range (μm) | −1.0 ; −2.0 | −1.0 ; −2.0 |
| Pixel size (Å) | 1.04 | 1.04 |
| Symmetry imposed | C1 | C1 |
| Micrographs (no.) | 6,544 | 6,544 |
| Total of extracted particles (no.) | 1,203,041 | 571,014 |
| Total of refined particles (no.) | 66,205 | 114,600 |
| Local resolution range (Å) | 3.26151 – 11.3835 | 3.54973 – 7.54117 |
| Resolution Masked 0.143 FSC (Å) | 3.46 | 3.69 |
| Refinement | 139 | 130 |
| Map sharpening B-factor (Å<sup>2</sup>) | −66 | −70 |
| Initial model used (PDB code) | 6yvu, 6yvd | 5oqn |
| Refinement package | Phenix (real space) | Phenix (real space) |
| r.m.s. deviations |  |  |
| Bond lengths (Å) | 0.005 | 0.006 |
| Bond angles (°) | 0.803 | 0.941 |
| Validation |  |  |
| MolProbity score | 1.31 | 1.29 |
| All-atom clashscore | 3.8 | 4.47 |
| Rotamers outliers (%) | 0.23 | 0.48 |
| Cβ outliers (%) | 0 | 0 |
| CaBLAM outliers (%) | 1.52 | 1.91 |
| B-factors (min / max / mean) |  |  |
| Protein | 21.5 / 135.51 / 70.04 | 56.22 / 127.96 / 81.81 |
| Nucleotides | 86.07 / 247.79 / 155.67 | 122.21 / 298.36 / 177.9 |
| Ligand | 33.54 / 64.65 / 59.39 | – |
| Overall correlation coefficients |  |  |
| CC (mask) | 0.81 | 0.76 |
| CC (peaks) | 0.71 | 0.7 |
| CC (volume) | 0.8 | 0.75 |
| Ramachandran plot statistics |  |  |
| Favored (%) | 97.2 | 97.68 |
| Allowed (%) | 2.8 | 2.32 |
| Disallowed (%) | 0 | 0 |
| EMRinger score | 2.55 | 1.28 |

**Table S2.**
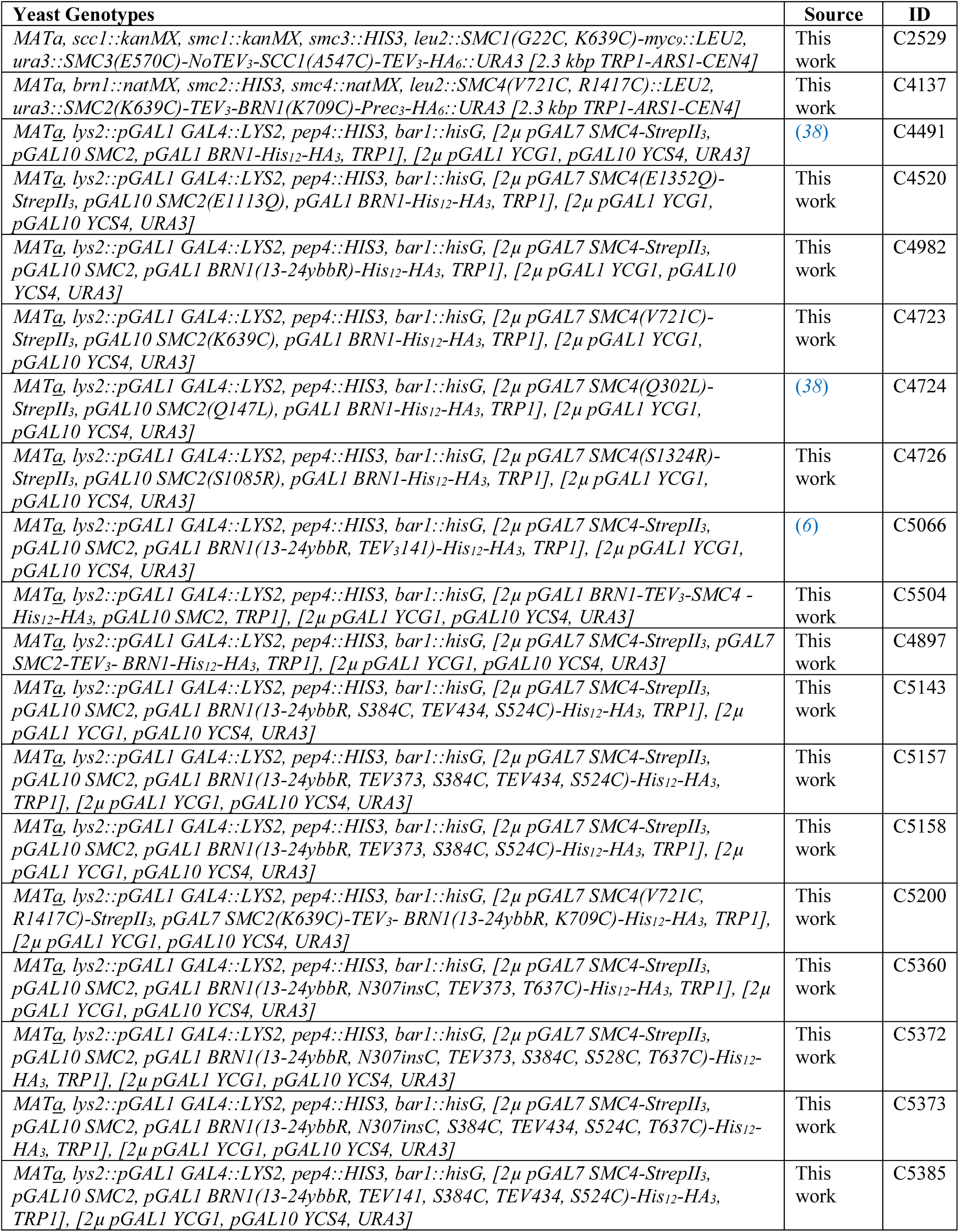
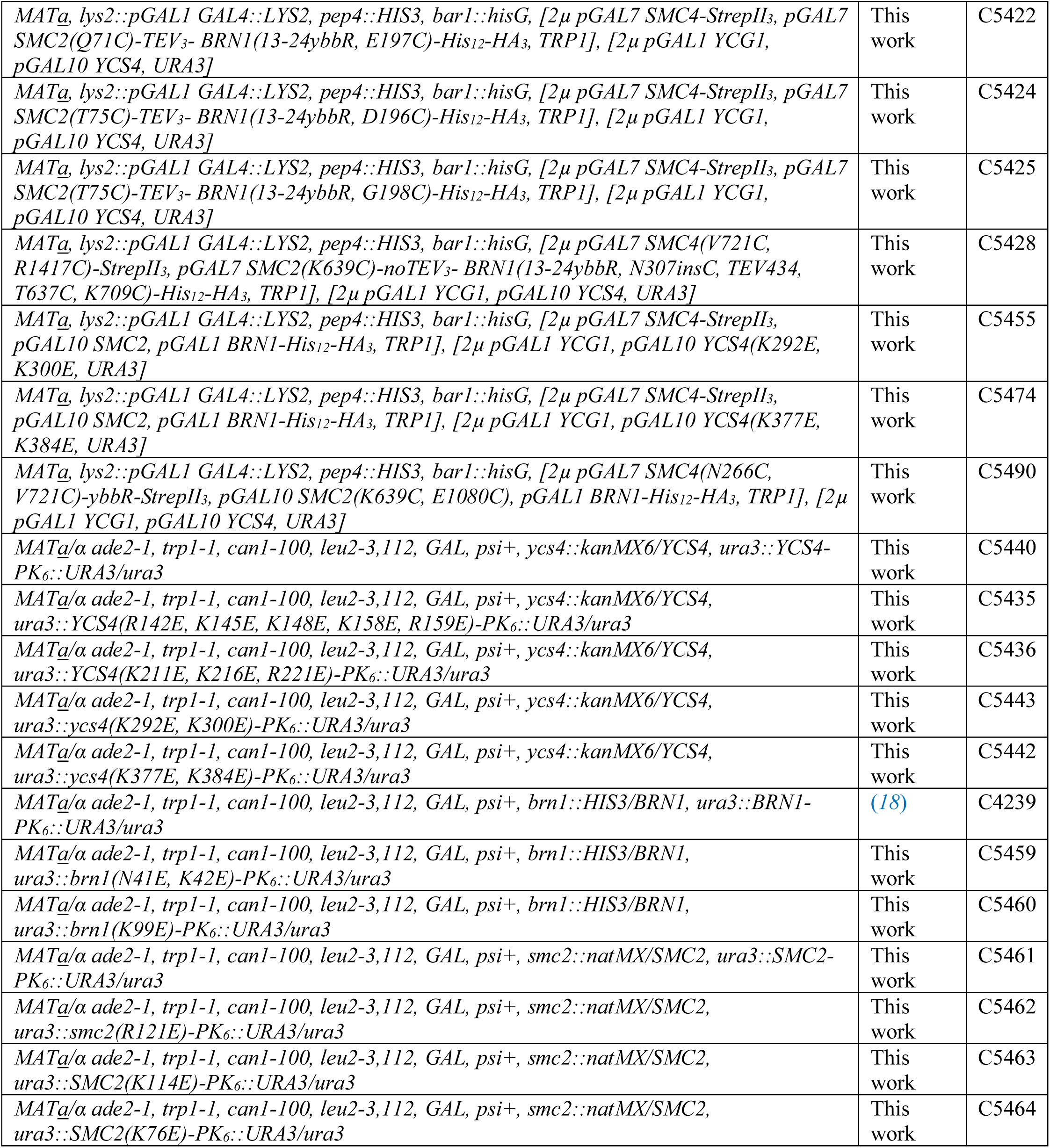
*Saccharomyces cerevisiae* strains.

**Movie S1. Fusion of the SMC-kleisin interfaces does not prevent DNA loop extrusion**. DNA loop extrusion DNA loop extrusion on SxO-stained double-tethered λ-phage DNA molecules by unmodified *Sc* condensin (top) or with Smc2–Brn1 (middle) or Brn1–Smc4 (bottom) fusions. Graphs plot normalized DNA intensities in, below and above the loop. Arrowheads mark the detected intensity maximum for segmentation of the loop.

**Movie S2. Animation of structural transitions between the apo and ATP/DNA-bound condensin core subcomplex**. Cartoon representation starting from the apo structure.

**Movie S3. Spontaneous rupture of DNA loops generated by non-cleavable condensin**. Loop formation and rupture on double-tethered λ-phage DNA molecules stained by SxO (in magenta or pseudo colors) by ATTO647N-labeled non-cleavable condensin (in cyan) in the presence of TEV protease. Positions of condensin at initial DNA binding (*t* = 0 s; grey arrowhead) and at loop rupture (white arrowhead) are indicated. Images are scaled to 4 × 10 µm.

**Movie S4. Rupture of DNA loops after TEV cleavage of kleisin chamber I**. As in movie S3 for a DNA loop generated by condensin with Brn1_TEV141_.

**Movie S5. Rupture of DNA loops after TEV cleavage of kleisin chamber II**. As in movie S3 for a DNA loop generated by condensin with Brn1_TEV434_.

**Movie S6. DNA loop extrusion by *Ct* holo condensin**. Unidirectional DNA loop formation on double-tethered λ-phage DNA by *Ct* holo condensin as in movie S1. DNA is stained with SxO (pseudo colors). The extrusion event lasts from 12 s to 22 s.

**Movie S7. DNA loop extrusion by *Ct* ΔYcg1 condensin**. DNA loop formation on double-tethered λ-phage DNA by *Ct* ΔYcg1 condensin as in movie S1. DNA is stained with SxO (pseudo colors). Loop formation starts at ∼2 s. The loop changes direction only once.

**Movie S8. DNA loop extrusion by *Ct* ΔYcg1 condensin**. DNA loop formation on double-tethered λ-phage DNA by *Ct* ΔYcg1 condensin as in movie S1. DNA is stained with SxO (pseudo colors). Loop formation starts at ∼14 s. The loop changes direction six times over a period of 140 s.

## References

1. I. F. Davidson, J. M. Peters, Genome folding through loop extrusion by SMC complexes. Nat Rev Mol Cell Biol, (2021).

2. S. Yatskevich, J. Rhodes, K. Nasmyth, Organization of Chromosomal DNA by SMC Complexes. Annu Rev Genet 53, 445–482 (2019).

3. M. S. van Ruiten, B. D. Rowland, On the choreography of genome folding: A grand pas de deux of cohesin and CTCF. Curr Opin Cell Biol 70, 84–90 (2021).

4. T. Hirano, Capturing condensin in chromosomes. Nat Genet 49, 1419–1420 (2017).

5. J. H. Gibcus et al., A pathway for mitotic chromosome formation. Science 359, (2018).

6. M. Ganji et al., Real-time imaging of DNA loop extrusion by condensin. Science 360, 102–105 (2018).

7. I. F. Davidson et al., DNA loop extrusion by human cohesin. Science 366, 1338–1345 (2019).

8. Y. Kim, Z. Shi, H. Zhang, I. J. Finkelstein, H. Yu, Human cohesin compacts DNA by loop extrusion. Science 366, 1345–1349 (2019).

9. S. Golfier, T. Quail, H. Kimura, J. Brugues, Cohesin and condensin extrude DNA loops in a cell cycle-dependent manner. Elife 9, (2020).

10. C. H. Haering, J. Lowe, A. Hochwagen, K. Nasmyth, Molecular architecture of SMC proteins and the yeast cohesin complex. Mol Cell 9, 773–788 (2002).

11. I. Onn, N. Aono, M. Hirano, T. Hirano, Reconstitution and subunit geometry of human condensin complexes. EMBO J 26, 1024–1034 (2007).

12. M. Kschonsak et al., Structural Basis for a Safety-Belt Mechanism That Anchors Condensin to Chromosomes. Cell 171, 588–600 e524 (2017).

13. Y. Li et al., Structural basis for Scc3-dependent cohesin recruitment to chromatin. Elife 7, (2018).

14. I. Piazza et al., Association of condensin with chromosomes depends on DNA binding by its HEAT-repeat subunits. Nat Struct Mol Biol 21, 560–568 (2014).

15. Z. Shi, H. Gao, X. C. Bai, H. Yu, Cryo-EM structure of the human cohesin-NIPBL-DNA complex. Science 368, 1454–1459 (2020).

16. T. L. Higashi et al., A Structure-Based Mechanism for DNA Entry into the Cohesin Ring. Mol Cell 79, 917–933 e919 (2020).

17. J. E. Collier et al., Transport of DNA within cohesin involves clamping on top of engaged heads by Scc2 and entrapment within the ring by Scc3. Elife 9, (2020).

18. M. Hassler et al., Structural Basis of an Asymmetric Condensin ATPase Cycle. Mol Cell 74, 1175–1188 e1179 (2019).

19. T. R. Beattie, S. D. Bell, Molecular machines in archaeal DNA replication. Curr Opin Chem Biol 15, 614–619 (2011).

20. M. H. Lamers et al., The crystal structure of DNA mismatch repair protein MutS binding to a G x T mismatch. Nature 407, 711–717 (2000).

21. L. Kashammer et al., Mechanism of DNA End Sensing and Processing by the Mre11-Rad50 Complex. Mol Cell 76, 382–394 e386 (2019).

22. G. Hauk, J. M. Berger, The role of ATP-dependent machines in regulating genome topology. Curr Opin Struct Biol 36, 85–96 (2016).

23. T. H. Massey, C. P. Mercogliano, J. Yates, D. J. Sherratt, J. Lowe, Double-stranded DNA translocation: structure and mechanism of hexameric FtsK. Mol Cell 23, 457–469 (2006).

24. T. G. Gligoris et al., Closing the cohesin ring: structure and function of its Smc3-kleisin interface. Science 346, 963–967 (2014).

25. C. H. Haering, A. M. Farcas, P. Arumugam, J. Metson, K. Nasmyth, The cohesin ring concatenates sister DNA molecules. Nature 454, 297–301 (2008).

26. M. Srinivasan et al., The Cohesin Ring Uses Its Hinge to Organize DNA Using Non-topological as well as Topological Mechanisms. Cell 173, 1508–1519 e1518 (2018).

27. S. Cuylen, J. Metz, C. H. Haering, Condensin structures chromosomal DNA through topological links. Nat Struct Mol Biol 18, 894–901 (2011).

28. B. G. Lee et al., Cryo-EM structures of holo condensin reveal a subunit flip-flop mechanism. Nat Struct Mol Biol 27, 743–751 (2020).

29. K. Hara et al., Structural basis of HEAT-kleisin interactions in the human condensin I subcomplex. EMBO Rep 20, (2019).

30. E. Kim, J. Kerssemakers, I. A. Shaltiel, C. H. Haering, C. Dekker, DNA-loop extruding condensin complexes can traverse one another. Nature 579, 438–442 (2020).

31. M. Hassler, I. A. Shaltiel, C. H. Haering, Towards a Unified Model of SMC Complex Function. Curr Biol 28, R1266–R1281 (2018).

32. J. F. Marko, P. De Los Rios, A. Barducci, S. Gruber, DNA-segment-capture model for loop extrusion by structural maintenance of chromosome (SMC) protein complexes. Nucleic Acids Res 47, 6956–6972 (2019).

33. M. L. Diebold-Durand et al., Structure of Full-Length SMC and Rearrangements Required for Chromosome Organization. Mol Cell 67, 334–347 e335 (2017).

34. F. Burmann et al., A folded conformation of MukBEF and cohesin. Nat Struct Mol Biol 26, 227–236 (2019).

35. B. Pradhan et al., SMC complexes can traverse physical roadblocks bigger than their ring size. bioRxiv, 2021.2007.2015.452501 (2021).

36. T. L. Higashi, G. Pobegalov, M. Tang, M. I. Molodtsov, F. Uhlmann, A Brownian ratchet model for DNA loop extrusion by the cohesin complex. Elife 10, (2021).

37. Y. Li et al., The structural basis for cohesin-CTCF-anchored loops. Nature 578, 472–476 (2020).

38. T. Terakawa et al., The condensin complex is a mechanochemical motor that translocates along DNA. Science 358, 672–676 (2017).

39. D. J. Fitzgerald et al., Protein complex expression by using multigene baculoviral vectors. Nat Methods 3, 1021–1032 (2006).

40. J. Schindelin et al., Fiji: an open-source platform for biological-image analysis. Nat Methods 9, 676–682 (2012).

41. M. Schorb, I. Haberbosch, W. J. H. Hagen, Y. Schwab, D. N. Mastronarde, Software tools for automated transmission electron microscopy. Nat Methods 16, 471–477 (2019).

42. J. Zivanov et al., New tools for automated high-resolution cryo-EM structure determination in RELION-3. Elife 7, (2018).

43. A. Punjani, J. L. Rubinstein, D. J. Fleet, M. A. Brubaker, cryoSPARC: algorithms for rapid unsupervised cryo-EM structure determination. Nat Methods 14, 290–296 (2017).

44. S. Q. Zheng et al., MotionCor2: anisotropic correction of beam-induced motion for improved cryo-electron microscopy. Nat Methods 14, 331–332 (2017).

45. A. Rohou, N. Grigorieff, CTFFIND4: Fast and accurate defocus estimation from electron micrographs. J Struct Biol 192, 216–221 (2015).

46. D. Tegunov, P. Cramer, Real-time cryo-electron microscopy data preprocessing with Warp. Nat Methods 16, 1146–1152 (2019).

47. T. Bepler et al., Positive-unlabeled convolutional neural networks for particle picking in cryo-electron micrographs. Nat Methods 16, 1153–1160 (2019).

48. P. B. Rosenthal, R. Henderson, Optimal determination of particle orientation, absolute hand, and contrast loss in single-particle electron cryomicroscopy. J Mol Biol 333, 721–745 (2003).

49. E. F. Pettersen et al., UCSF ChimeraX: Structure visualization for researchers, educators, and developers. Protein Sci 30, 70–82 (2021).

50. P. Emsley, B. Lohkamp, W. G. Scott, K. Cowtan, Features and development of Coot. Acta Crystallogr D Biol Crystallogr 66, 486–501 (2010).

51. J. Jumper et al., Highly accurate protein structure prediction with AlphaFold. Nature 596, 583–589 (2021).

52. T. I. Croll, ISOLDE: a physically realistic environment for model building into low-resolution electron-density maps. Acta Crystallogr D Struct Biol 74, 519–530 (2018).

53. P. V. Afonine et al., Real-space refinement in PHENIX for cryo-EM and crystallography. Acta Crystallogr D Struct Biol 74, 531–544 (2018).

54. J. F. Marko, E. D. Siggia, Statistical mechanics of supercoiled DNA. Phys Rev E Stat Phys Plasmas Fluids Relat Interdiscip Topics 52, 2912–2938 (1995).

